# Metagenomics of Parkinson’s disease implicates the gut microbiome in multiple disease mechanisms

**DOI:** 10.1101/2022.06.08.495316

**Authors:** Zachary D Wallen, Ayse Demirkan, Guy Twa, Gwendolyn Cohen, Marissa N Dean, David G Standaert, Timothy Sampson, Haydeh Payami

## Abstract

Parkinson’s disease (PD) may start in the gut and spread to the brain. To investigate the role of gut microbiome, we enrolled 490 PD and 234 control individuals, conducted deep shotgun sequencing of fecal DNA, followed by metagenome-wide association studies requiring significance by two methods (ANCOM-BC and MaAsLin2) to declare disease association. Thirty-percent of species and pathways tested had altered abundances in PD, depicting a widespread dysbiosis. Network analysis showed PD-associated species form polymicrobial clusters that grow or shrink together, and some compete. Metagenomic profile of PD indicates a disease permissive microbiome, evidenced by overabundance of pathogens and immunogenic components, dysregulated neuroactive signaling, preponderance of molecules that induce alpha-synuclein pathology, and over-production of toxicants; with the reduction in anti-inflammatory and neuroprotective factors limiting the capacity to recover. These data provide a broad foundation with a wealth of concrete testable hypotheses to discern the role of the gut microbiome in PD.

## Introduction

Microbiota (the billions of microorganisms living inside and on the human body) are necessary for human health. The gut microbiome (the collective genomes of microbiota that inhabit human gut) aid with dietary metabolism, produce essential metabolites such as vitamins, maintain the integrity of the intestinal barrier, inhibit pathogens, and metabolize drugs and toxicants^1^. The gut microbiome regulates development and continued education of the host’s immune response and nervous system by producing specific metabolites with far reaching effects such as the maturation and maintenance of microglia in the brain^2^. An imbalance (dysbiosis) in microbiome composition and function can render host prone to disease. Studies in humans and animal models have revealed disease-related dysbiosis in a range of common metabolic (e.g., diabetes), inflammatory (inflammatory bowel disease), neurologic (Parkinson’s disease) and developmental disorders (autism)^3^.

Parkinson’s disease (PD) is a progressively debilitating disorder that affected 4 million Individuals in year 2005 and is projected to double to 8.7 million individuals by year 2030. Although historically defined as a movement disorder, PD is a multi-systemic disease^4^. The earliest sign is often constipation which can precede motor signs by decades^5^. Moreover, PD is etiologically heterogenous. Despite the discovery of several causative genes^6^, 90 susceptibility loci spanning the human genome^7^, and multiple environmental risk factors^8^, the vast majority of PD remains idiopathic. It is speculated that PD is caused by various combinations of genetic susceptibility and environmental triggers, although no causative combination has yet been identified.

Braak’s hypothesis^9^ that non-familial forms of PD start in the gut by a pathogen is gaining increasing support. The connection between PD and the gastrointestinal (GI) system, including constipation, compromised gut barrier, and inflammation, has long been established. Alpha-synuclein pathology has been detected in the gut of persons with PD at early stages^10^, and there is evidence from imaging studies that in some cases pathology may start in the gut and spread to the brain^11^. In mice, it was shown that alpha-synuclein fibrils injected into gut induce alpha-synuclein pathology which spreads from gut to brain, and that vagotomy stops the spread^12^. In parallel, large epidemiological studies have shown that persons who had complete truncal vagotomy decades earlier had substantially reduced incidence of PD later in life^13,14^.

With accumulating evidence implicating the gut as an origin of PD, and the newly gained appreciation for the involvement of gut microbiome in chronic diseases, there has been increasing interest in decoding the connection between the gut microbiome and PD. In mice overexpressing the human alpha-synuclein gene, we have shown that gut microbiome regulates alpha-synuclein mediated pathophysiologies^15^. In another genetic model of PD (*Pink1*^*-/-*^), intestinal infection with Gram-negative bacterial pathogens was shown to elicit an immune reaction that leads to neuronal degeneration and motor deficits, and which can be reversed with the PD-medication, L-DOPA^16^. In addition, we and others have found that, curli, an amyloidogenic protein produced by Gram-negative *Escherichia coli*, induces alpha-synuclein aggregation and accelerates disease in the gut and neurodegeneration in the brain^17-19^. We have detected overabundance of opportunistic pathogens in the gut microbiomes of individuals with PD^20^. Collectively, experimental and human studies support Braak’s hypothesis that intestinal infection may act as a triggering event in PD, but it is yet to be proven that pathogens in human gut cause PD.

Studies conducted on human fecal samples have all found evidence of dysbiosis in PD gut microbiome but results on specific microorganisms that drive the dysbiosis have been mixed^21-23^. Human studies of PD and microbiome have had limited sample sizes and nearly all were based on 16S rRNA gene amplicon sequencing (henceforth, 16S) which limits resolution to genus-level. Metagenomics (study of all genetic material sampled from a community) is an emerging field in medical science. The microbiome can now be studied in large-scale human studies at high resolution of species and genes via metagenomics. Here we present the first large-scale metagenomics analysis of PD gut microbiome. This study was designed and executed by a single team of investigators (NeuroGenetics Research Consortium, NGRC), enabling complete control to employ state of the art methods and ensure uniformity from start to end.

## Results and Discussion

### The cohort

As outlined in STORMS^24^ flowchart (Fig. 1), the study included a newly enrolled cohort of 490 persons with PD and 234 neurologically healthy controls (NHC). All subjects were from a single geographic area in deep-south United States (US), minimizing confounding by geographic variation. Fifty-five percent of NHC were spouses and shared environment with PD. Using uniform methods, we collected extensive metadata (Supplementary Fig. 1) and a stool sample from each subject, extracted DNA from stool and conducted deep shotgun sequencing achieving average 50M raw reads/sample. This dataset is new and is publicly available. The sample size is comparable to Human Microbiome Project (HMP) which included 242 healthy individuals age 18-40 years, 100 individuals with inflammatory bowel disease, and 106 individuals with pre-diabetes^25^. The older ages of the controls, and their neurologically healthy status, is a unique addition to the publicly available datasets.

**Fig. 1.**
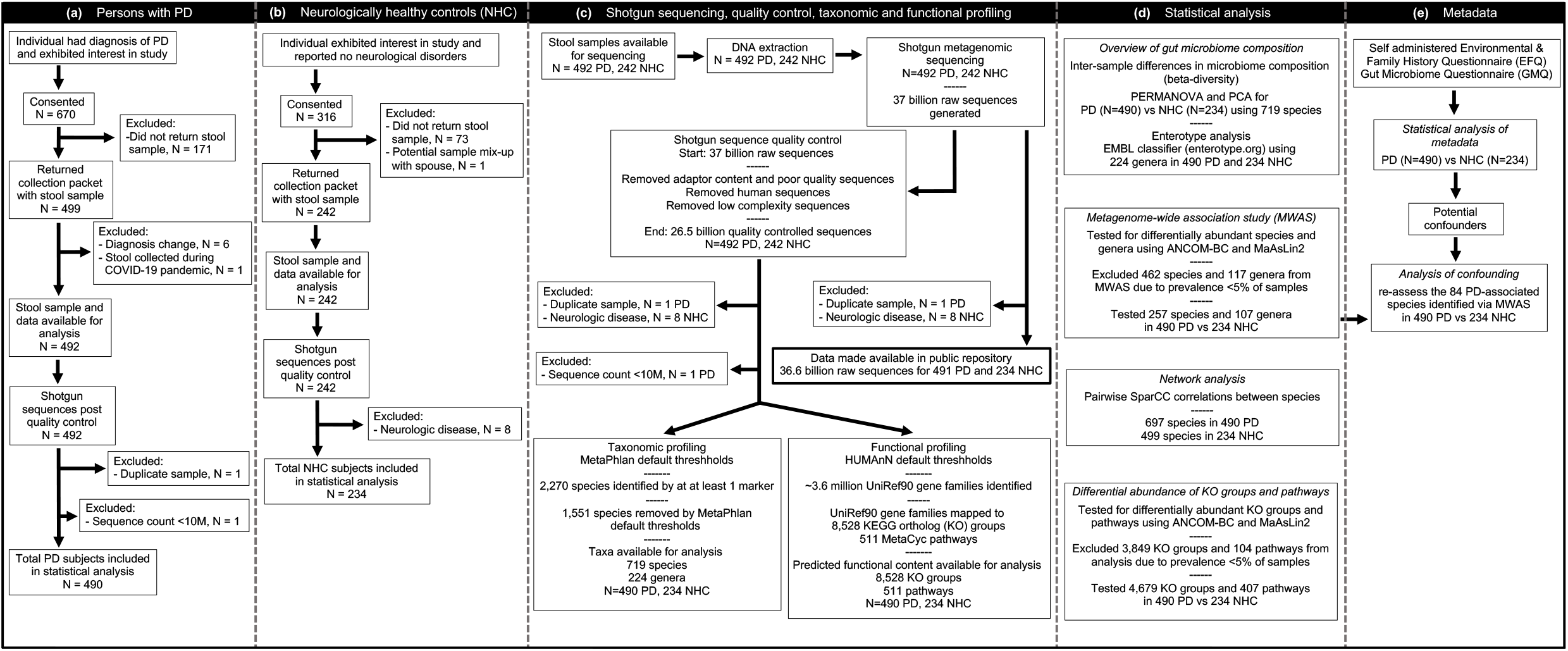
The STORMS flowchart. Following reporting guidelines for human microbiome research^24^.

### Metadata

Data on 53 variables were analyzed to characterize the subjects, and to identify disease-associated variables that could potentially confound downstream metagenomics analyses (Table 1). Gastrointestinal (GI) problems, which are well-known features of PD, were readily evident in this cohort. Constipation was more prevalent in PD cases (odds ratio (OR)=6.1, P=2E-19 for chronic constipation, P=3E-6 for Bristol Chart score), and PD cases reported more GI discomfort than NHC (OR=2.8, P=3E-7). Compared to NHC, PD cases had diminished intake of alcohol (OR=0.6, P=3E-4) and foods in all five categories (fruits/vegetables, animal products, nuts, yogurt, and grains), all reaching significance (OR=0.6-0.7, P=0.002-0.05) except grains (OR=0.8, P=0.2). Use of laxatives (OR=3.8, P=7E-10), pain medication (OR=1.6, P=0.04), sleep aid (OR=2, P=7E-5), and medication for depression/anxiety/mood (OR=2.1, P=7E-5) were more common in PD than NHC. Probiotic supplement use was more common in NHC than PD (OR=0.6, P=0.02), which is noteworthy because as the data will show, *Bifidobacterium* and *Lactobacillus* species, which are common constituents of commercial probiotics, were more abundant in PD than NHC metagenomes. Variables that differed in PD vs. NHC were evaluated as potential confounders in downstream metagenomics analyses.

**Table 1.**
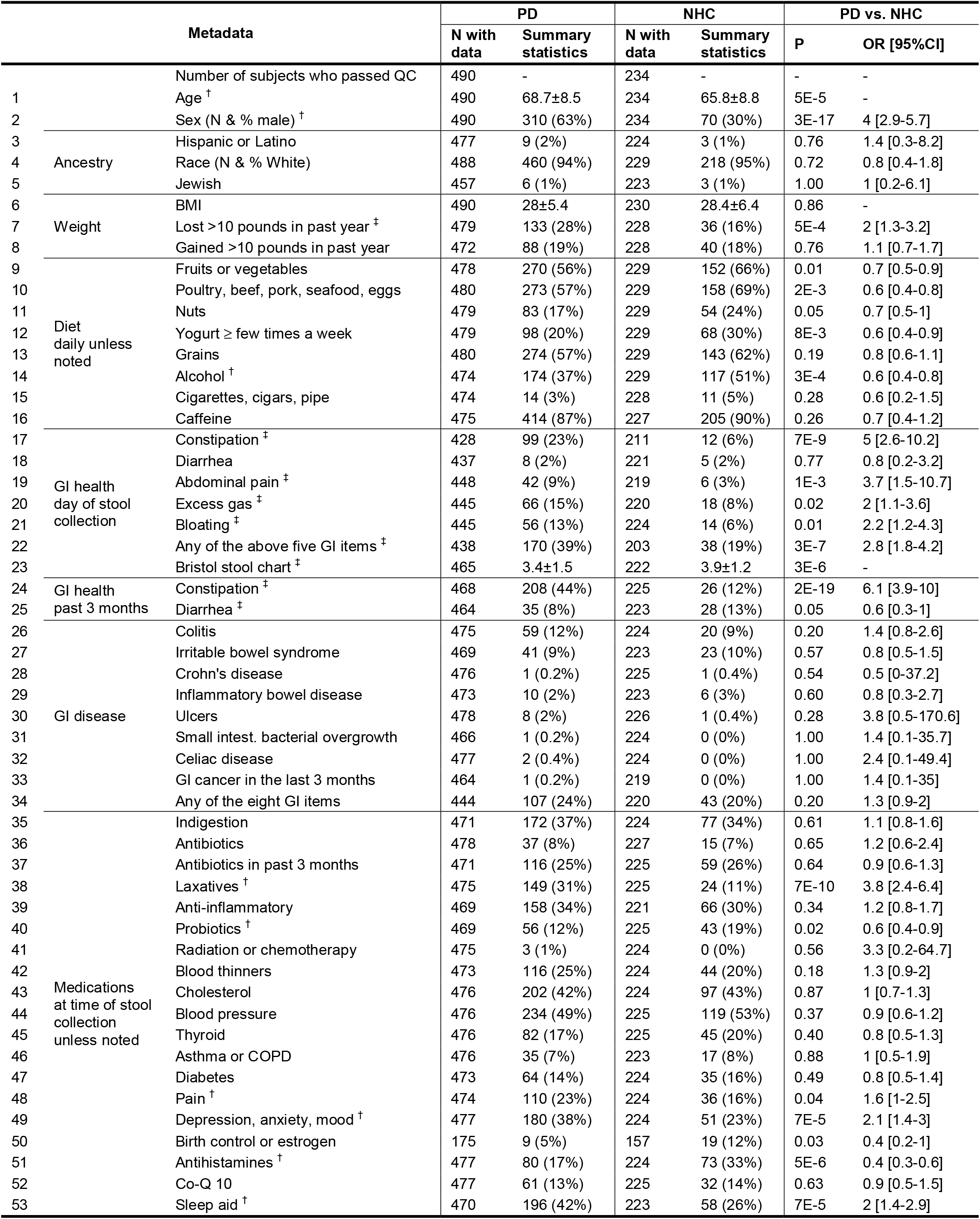
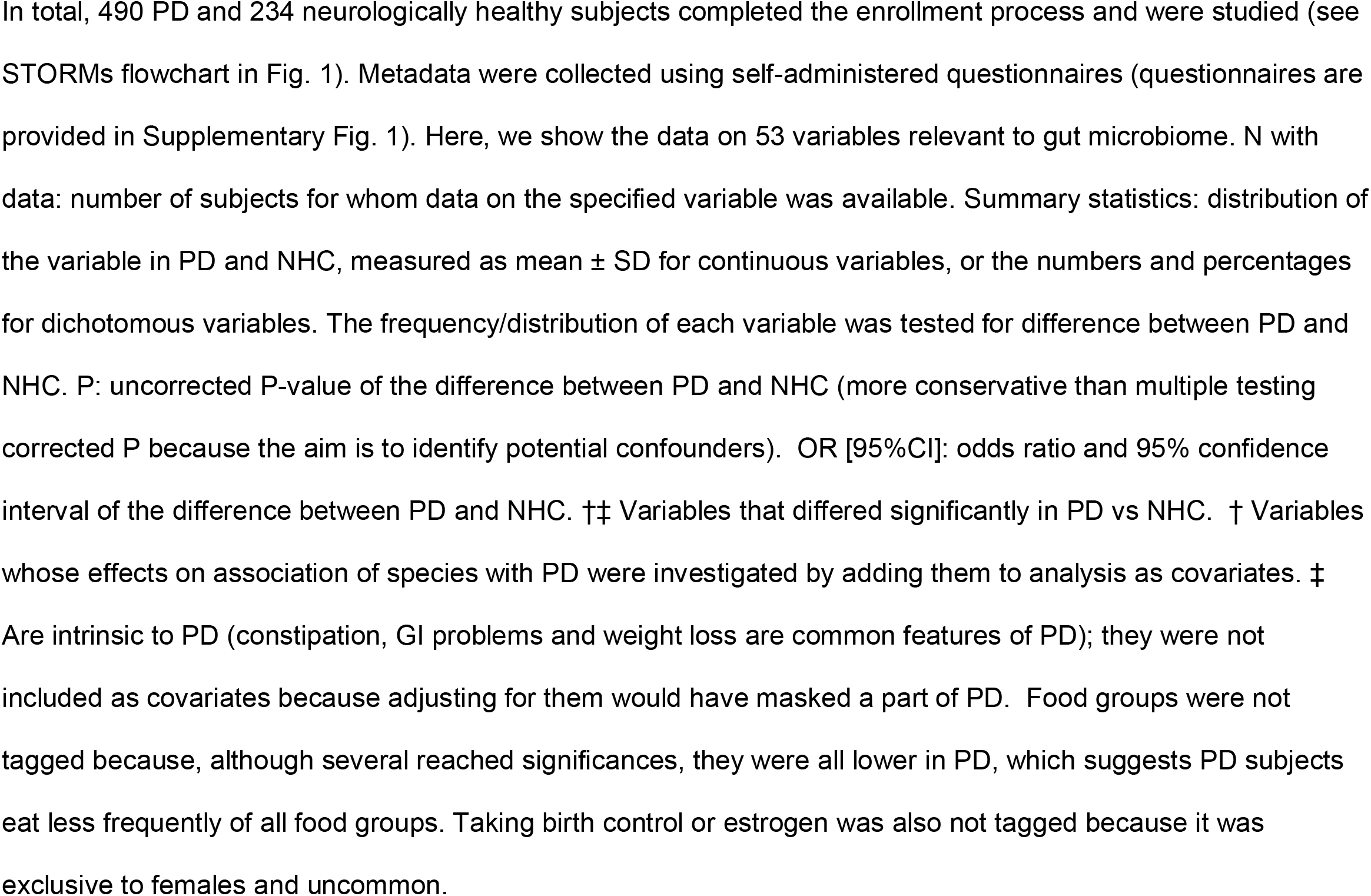
Subject characteristics and metadata.

### Profiles of PD and NHC metagenomes

Inter-individual difference in the global composition of the gut metagenome, beta-diversity, was significantly different in PD vs. NHC (P<1E-4). Enterotype profiles of PD were also significantly different from NHC (P=4E-4) driven primarily by enrichment of *Firmicutes* in PD samples (Supplementary Fig. 2).

A major goal of microbiome studies is to identify the disease-associated species. Including Bacteria, Archaea and Eukarya, we identified a total of 2,270 species, 719 of which passed stringent bioinformatic quality control (QC) thresholds, and 257 were present in >5% of subjects. We conducted an unbiased metagenome-wide association study (MWAS) to test differential abundances of species in PD vs. NHC, using two statistical methods (MaAsLin2 and ANCOM-BC). We also conducted MWAS at genus-level to be able to interpret our results in the context of the existing literature. The full MWAS results are provided in Supplementary Tables 1 and 2. We nominated a species or genus as PD-associated if it achieved significance by MaAsLin2 and ANCOM-BC (i.e., false discovery rate (FDR)<0.05 by one and FDR≤0.1 by the other). Of the 257 species tested, 84 were associated with PD: 55 were enriched and 29 were depleted in PD (Table 2, Supplementary Fig. 3). Of the 107 genera tested, 34 were associated with PD: 23 were enriched and 11 were depleted in PD (Table 3). Thus, whether measured at species or genus-level, the dysbiosis in the PD gut microbiome appears to involve about 30% of tested taxa.

**Table 2.**
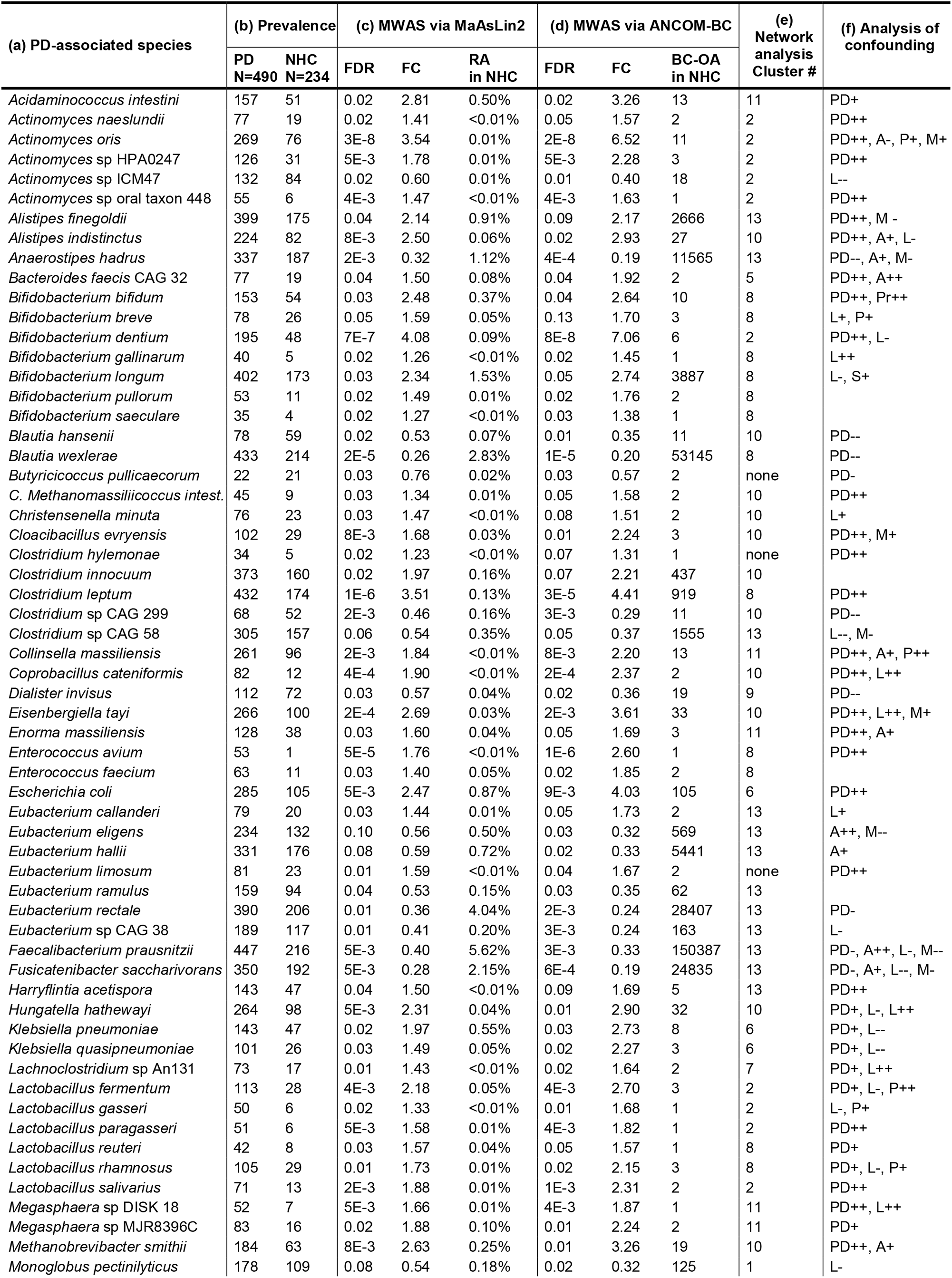
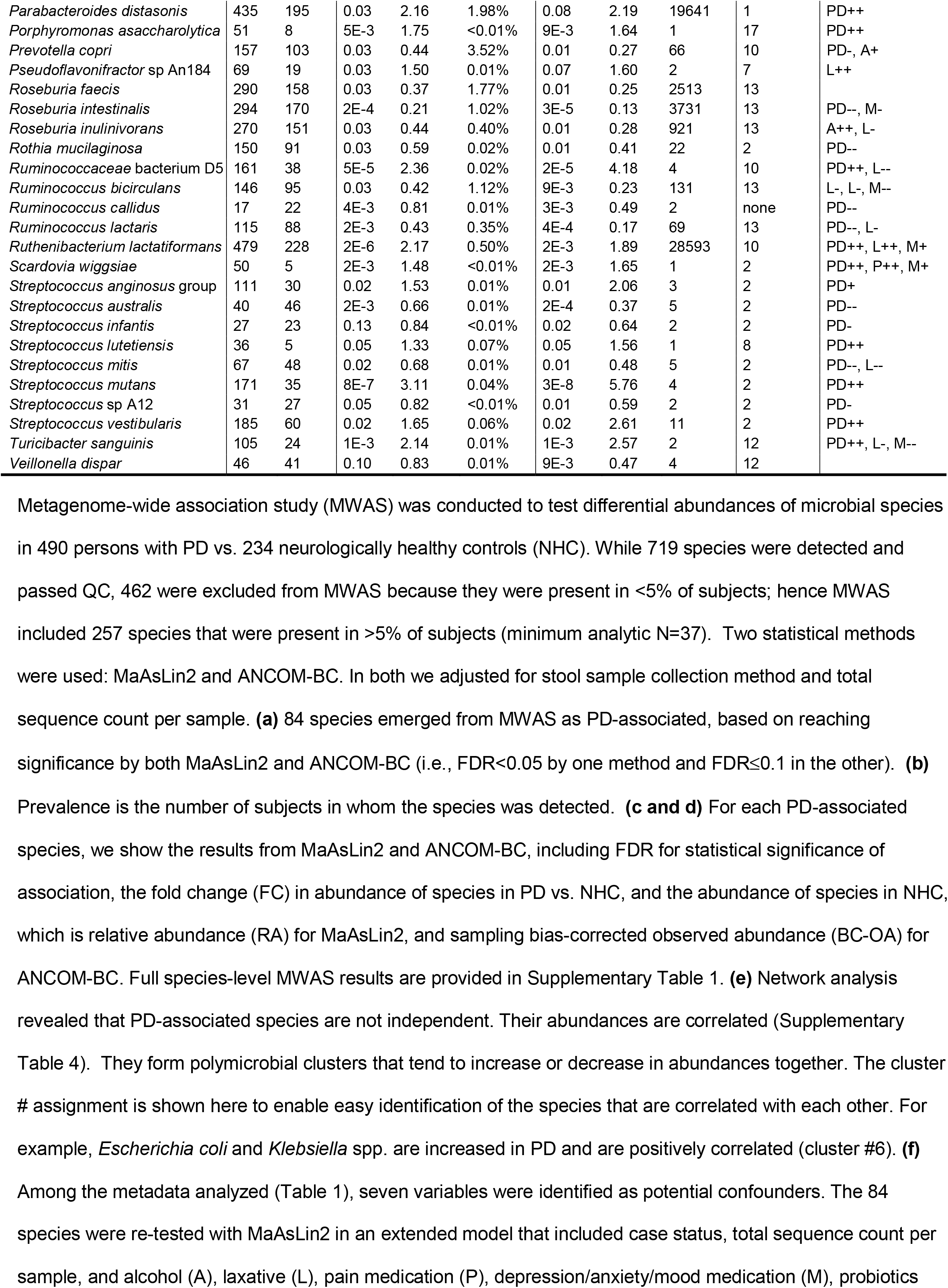
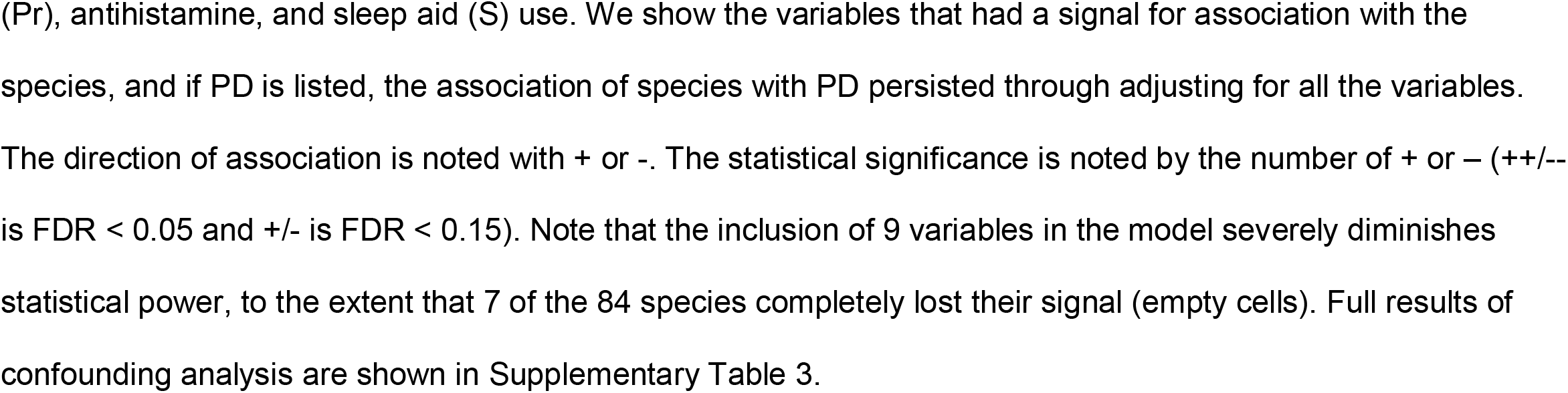
Identification and characterization of 84 PD-associated species.

**Table 3.**
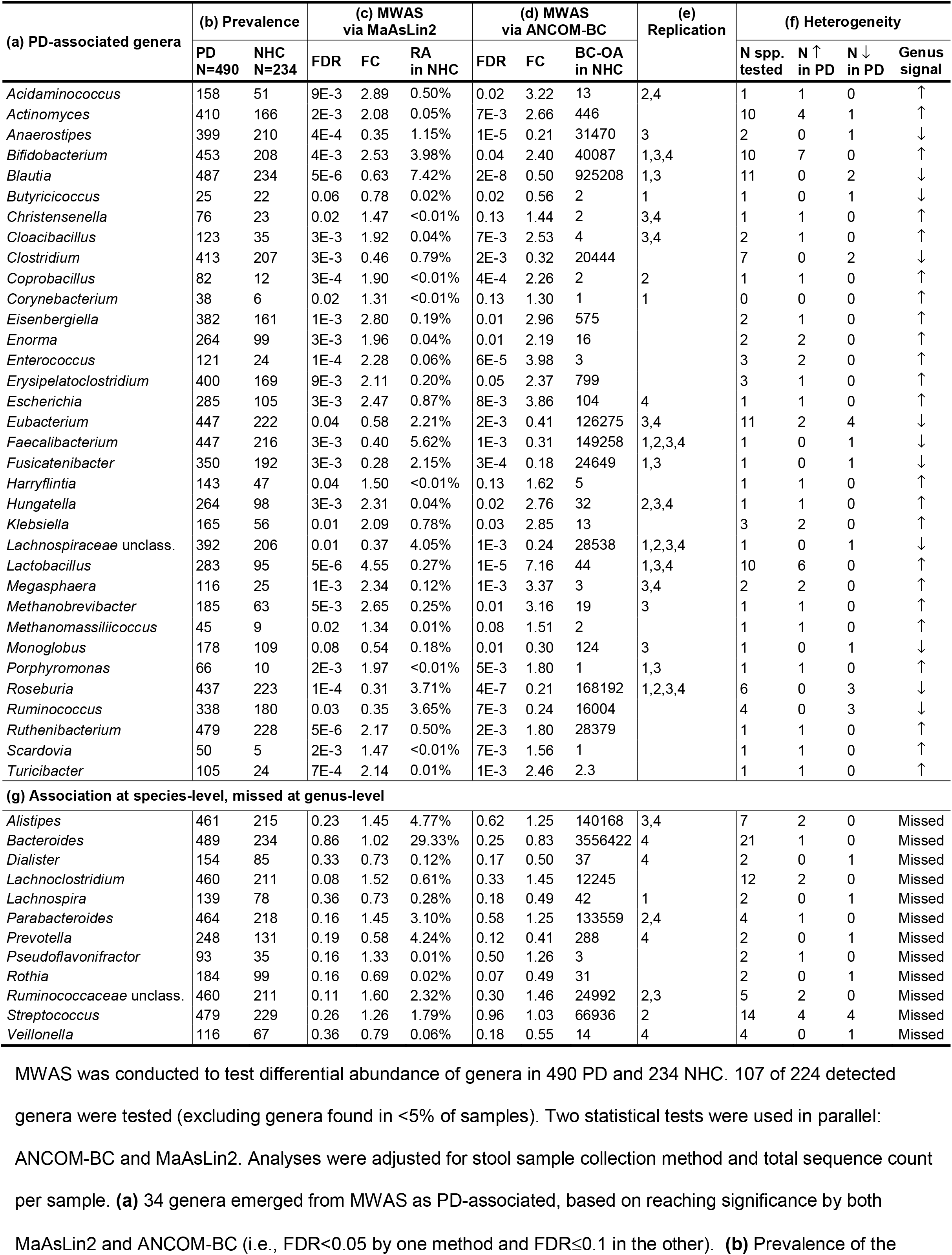
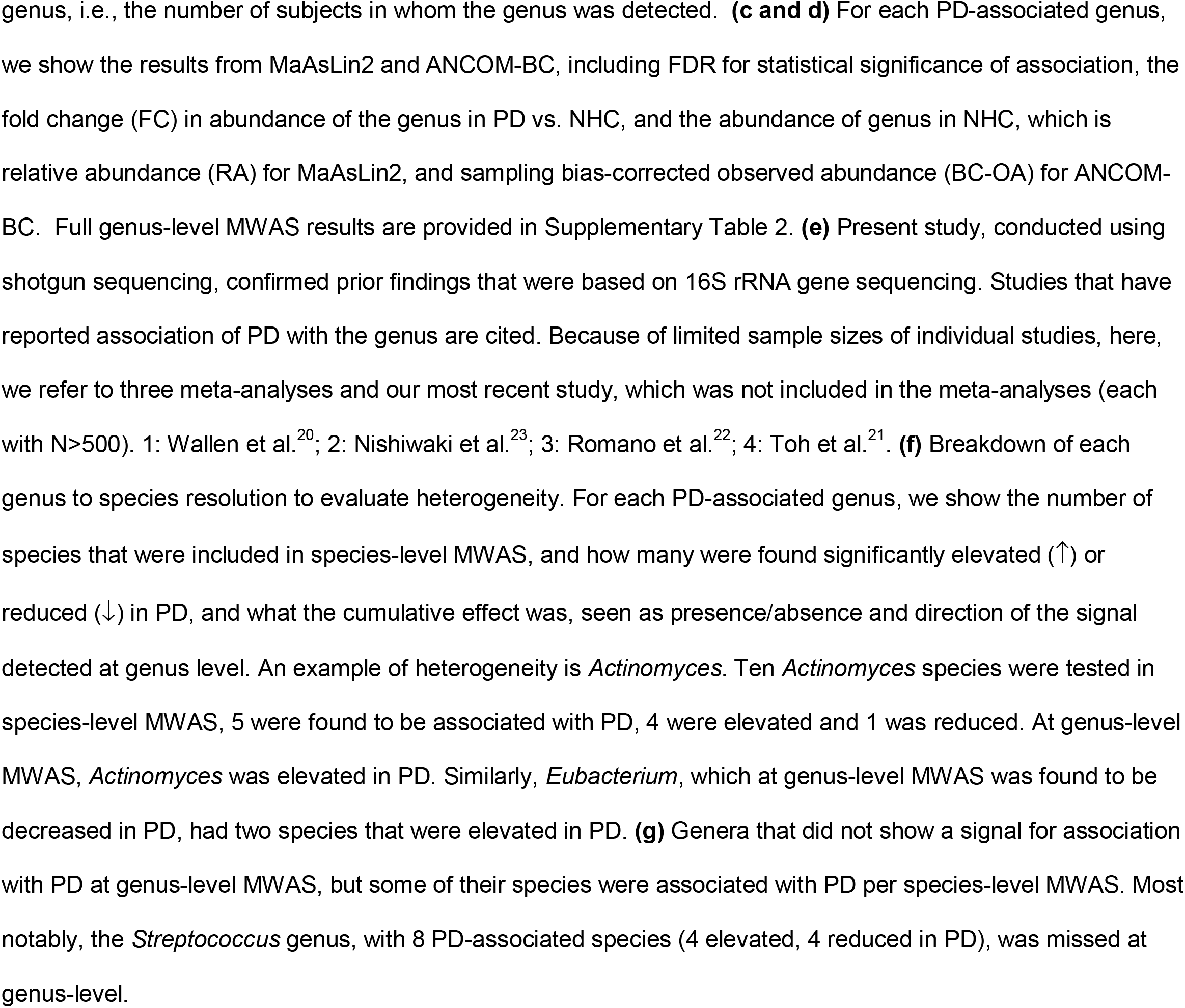
PD-associated genera, replication of prior findings, and evidence of heterogeneity.

The primary aim of this study was to generate a full, unaltered view of the dysbiosis in PD gut microbiome. To that end, MWAS and downstream analyses were adjusted only for technical variables (i.e., total sequence count per sample, and stool collection kit). Adjusting for variables like constipation, which are intrinsic to PD, would mask their effect in the full view that we are aiming for. However, as secondary analyses, we investigated effects of age and sex, and explored confounding by extrinsic variables that had differing frequencies/distributions in PD vs. NHC, namely intake of alcohol, laxative, probiotics, antihistamines, depression/anxiety/mood medication, pain medication, and sleep aid (Table 1). We tested association of the 84 species, that had emerged from MWAS, with PD while including age and sex in the model, and all species retained significance for association with PD at FDR<0.1. Next, we re-tested the 84 species in a model that included the extrinsic variables (Supplementary Table 3). Statistical power was substantially reduced when testing all variables simultaneously, to a degree that species that were rare and had smaller analytic sample sizes lost signal all together. Nonetheless, we confirmed the association of PD with 62 of 84 species (Table 2): 32 species were associated only with PD, while the other 30 were associated with PD and one or more other variables, most commonly alcohol (avoided by PD) or laxatives (frequently used by PD).

### Network analysis reveals clusters of co-occurring and competing species

We calculated pairwise correlation (r) in species abundances for PD metagenome using all 697 species detected (Supplementary Table 4) and NHC metagenome using 499 species detected (Supplementary Table 5). We observed both positive and negative correlations throughout the metagenome, including PD-associated species. Positive correlation among species suggests they tend to co-occur in the same sample and their abundances rise or fall together, while negative correlation suggests that, within a sample, an increase in the abundance of one species correlates with decrease in the abundance of another. We used pairwise correlations reaching |r| >0.2 and permutation P<0.05 to create correlation networks and algorithmically defined clusters of correlated species (Fig. 2a, Supplementary Fig. 4, Supplementary Table 6). We then mapped PD-associated species to the networks (Fig. 2b).

**Fig. 2.**
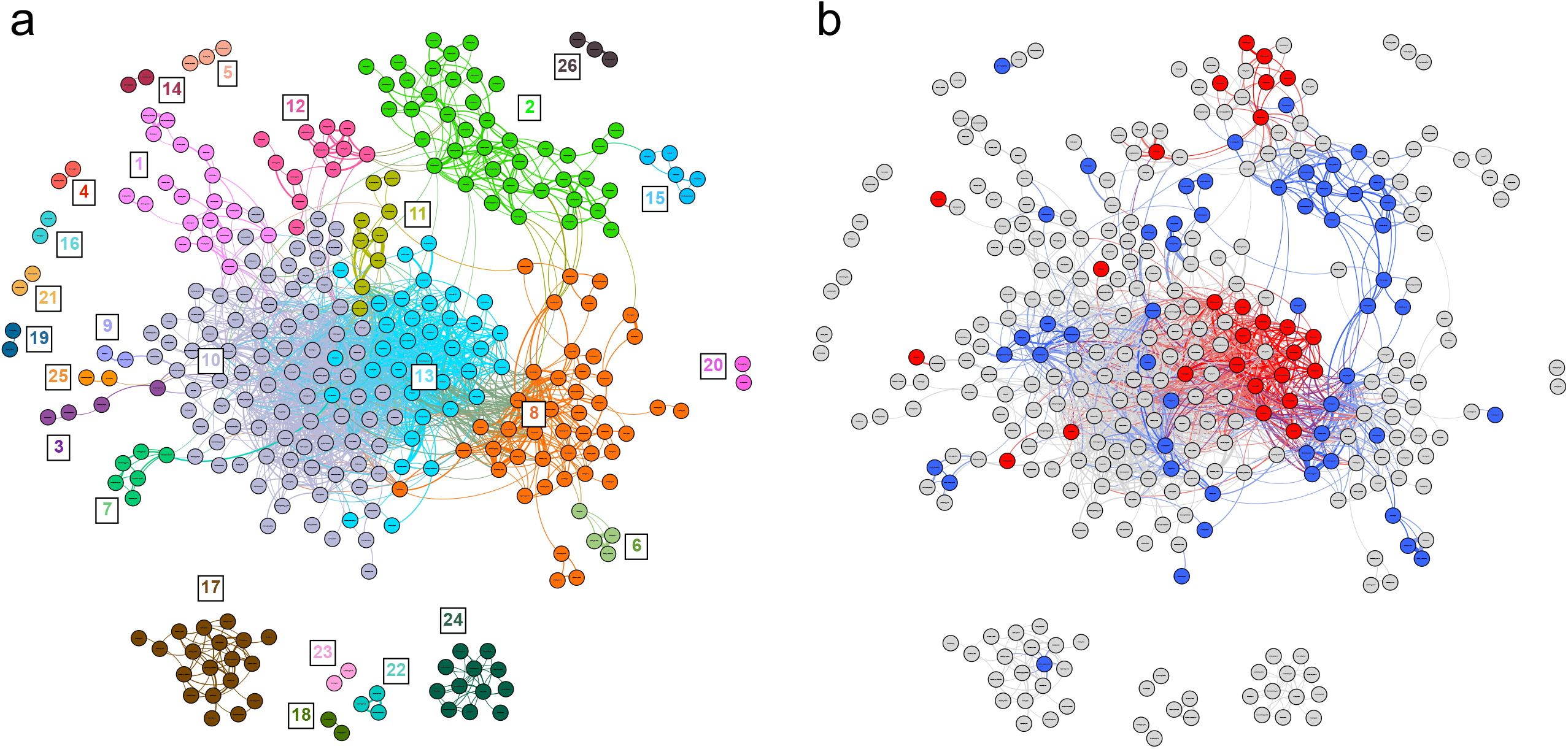
Network analysis reveals polymicrobial clusters of correlated species in the PD metagenome. **(a)** All species detected in PD gut metagenome were plotted if their abundance correlated with at least one other species (i.e., |r| > 0.2 and permuted P-value < 0.05). Clusters were defined by Louvain algorithm and were randomly assigned a color and a number. Each circle (node) denotes a species and the curved lines (edges) connect correlated species. **(b)** PD-associated species that were identified via MWAS were mapped to the network, and highlighted in blue if elevated in PD, or red if reduced in PD. Correlation among PD-associated species were often positive, indicating abundances tended to rise together like *Escherichia coli, Klebsiella pneumoniae*, and *Klebsiella quasipneumoniae* in cluster #6, or decline together like the polysaccharide metabolizing species in cluster #13. Sometimes the correlation was negative indicating increase in abundance of one species correlated with decrease in the other, e.g., in cluster #2 the rise in abundances of some *Streptococcus* species in PD microbiome correlated with a decline in the abundances of other *Streptococcus* species.

Species that showed the greatest increase in PD including *Bifidobacterium dentium, Actinomyces oris, Streptococcus mutans, Lactobacillus fermentum* and several other *Lactobacillus, Actinomyces*, and *Streptococcus* species were positively correlated with each other and mapped to cluster #2 of the PD network (Fig. 2). Cluster #2 included 41 species, 19 were associated with PD: 13 enriched, 6 reduced in PD, all connected via positive and negative correlations. Among them were 8 *Streptococcus* species: 4 elevated and 4 reduced in PD. Notably, *Streptococcus mutans* (elevated in PD) was negatively correlated with *Streptococcus sp. A12* (reduced in PD), which aligns with the original characterization of *Streptococcus sp. A12* as a novel strain that inhibits the growth and signaling pathways of the pathogenic *Streptococcus mutans*^26^.

Most of the species that were depleted in PD, including short chain fatty acid (SCFA) producing species of *Roseburia, Eubacterium, Ruminococcus*, and *Faecalibacterium prausnitzii* were positively correlated with each other and mapped to cluster #13 of the PD network (Fig. 2). Moreover, decreasing abundances of SCFA producing species were correlated with increasing abundances of *Bifidobacterium* species (Fig. 2b, Supplementary Table 4). These findings suggest competitive interactions occurring in the dysbiotic PD microbiome, both at large community scale (*Bifidobacterium* species vs. SCFA-producing species), as well as at species scale (*Streptococcus sp. A12* vs. *Streptococcus mutans*).

We noted two clusters with species that are known to cause infections. *Escherichia coli, Klebsiella pneumoniae* and *Klebsiella quasipneumoniae*, which were elevated in PD, were positively correlated and grouped into cluster #6 of the PD network. Cluster #17 is a polymicrobial community of 19 opportunistic pathogen species (Supplementary Table 7). We detected overabundance of these taxa at genus-level in our prior datasets^20^. Here, at species-level, only *Porphyromonas asaccharolytica* was included in MWAS and it confirmed to be significantly elevated in PD. The other 18 species were too rare individually to be tested in MWAS. When taken together, the relative abundance of species in cluster #17 was significantly elevated in PD (fold change (FC)=2.63, P=2E-5). Not only the prevalence and abundance of these species were elevated in PD, but they also formed a tightly interconnected polymicrobial cluster in PD metagenome, as they do in clinical infection specimen^27^, but not in NHC metagenome. In PD, each species in cluster #17 connects with an average of 5.4 other species, as compared to 1 in NHC (Supplementary Table 7).

### Replication

We have previously characterized two cohorts of PD and NHC subjects using 16S sequencing (N1=333 and N2=507)^20,28^. Except for 11 subjects who were inadvertently double-enrolled, the 724 subjects in this shotgun study are independent of the 840 subjects in the 16S studies. Having these three large datasets presented a unique opportunity to compare 16S and shotgun results, and replicate and validate the findings. In prior 16S studies^20,28^, we reported association of 15 genera with PD and showed that they form three clusters of (1) opportunistic pathogens (elevated in PD), (2) SCFA-producing bacteria (reduced), and (3) probiotics *Bifidobacterium* and *Lactobacillus* (elevated). Here, we readily confirmed 13 of the 15 genus-level associations with statistical significance (Table 3). The remaining two (*Oscillospira, Prevotella* sub-genus) were not captured here due to differences in reference databases used for taxonomic assignment: SILVA v132 for 16S and ChocoPhlAn for shotgun. *Oscillospira*, a genus designation in SILVA, was not in ChocoPhlAn. *Prevotella*, as called by SILVA, was in fact a sub-genus of *Prevotella* and included species *P. buccalis, P. timonensis, P. disiens, P. bivia, P. amnii*, and *P. oralis*. Although detected, none of these species were tested because they were individually rare. We grouped them to recreate the *Prevotella* sub-genus, and successfully replicated the positive association with PD (FC=1.5, P=1E-3). We also captured and replicated the previously defined genus clusters of opportunistic pathogens (cluster #17 here), SCFA-producing bacteria (cluster #13), and “probiotics” *Bifidobacterium* and *Lactobacillus* (cluster #2). Thus, using metagenomics, we replicated both MWAS and network analysis findings of 16S studies in an independent dataset, and resolved them to species-level.

### Alignment with existing literature

Prior studies of PD and microbiome were conducted using 16S, and while all detected a significant dysbiosis in PD gut, the results on PD-associated taxa were widely inconsistent. Small sample sizes and inter-study variations were thought responsible. Recent meta-analyses were able to identifiy few associations at family- or genus-level robust enough to transcend inter-study variations^21-23^. We successfully confirmed these associations at genus-level and resolved them to species, including *Blautia, Faecalibacterium, Fusicatenibacter, Roseburia* and *Ruminococcus*, which are reduced in PD, and *Bifidobacterium, Hungatella, Lactobacillus, Methanobrevibacter* and *Porphyromonas*, which are elevated in PD^21-23^. *Prevotella* has been reported as decreased in PD by some^29,30^ and increased by others^20,28^. At species-level, *Prevotella copri* was decreased, and the pathogenic species of *Prevotella* (as defined by SILVA above) were increased as a group, confirming, and resolving the seemingly contradictory reports on *Prevotella. Akkermansia* is a conundrum. Most studies report elevated *Akkermansia* in PD. We did not detect a statistically significant signal for *Akkermansia* at genus- or species-level in this southern US cohort, although the trend was elevated. Interestingly, *Akkermansia* was elevated in our two prior 16S datasets as well, but reached significance only in the multi-state cohort that was primarily from northern US^28^ and not the dataset from southern US^20^, suggesting a geographic effect.

Analyses conducted at genus or higher taxonomic levels have the underlying assumption that microorganisms within each classification have similar trend of association with disease and can therefore be collapsed. If the assumption does not hold, signals may be missed or be inconsistent across studies (as was shown for *Prevotella* above). We encountered several such heterogenous genera (Table 3). Most notably, we detected 8 PD-associated *Streptococcus* species: 4 were elevated in PD, 4 were reduced. However, at genus-level, *Streptococcus* lacked evidence for association with PD. Thus, *Streptococcus*, the genus with the greatest number of PD-associated species, was missed at genus-level because of heterogeneity.

### Complementarity of species-, genus- and cluster-level analyses

Species-level testing can capture the signals that are lost at genus-level due heterogeneity within genus. In fact, we detected associations with 22 species belonging to 12 genera where the associations were missed at genus-level MWAS (Table 3). Conversely, when species are rare and excluded from testing for statistical reasons, the signal may be present at genus-level due to cumulative effects of species with similar effects (e.g., *Corynebacterium*). Networks of correlating species is another complimentary source of information as it may reveal disease relevant polymicrobial clusters. Consider cluster #17 composed of 19 opportunistic pathogen species from 10 genera: species-level MWAS detected only one of the 19 species as elevated in PD (*Porphyromonas asaccharolytica*), genus-level MWAS detected 3 of the 10 genera as elevated in PD (*Actinomyces, Corynebacterium, Porphyromonas*), and cluster-level analysis detected the whole polymicrobial cluster as elevated in PD.

### Microbial gene-families and metabolic pathways

We identified 8,528 gene-families (KEGG ortholog (KO) groups) and 511 metabolic pathways. Excluding those present in <5% of subjects, 4,679 KO groups and 407 metabolic pathways were tested for differential abundance in PD vs. NHC using MaAsLin2 and ANCOM-BC. The full results are provided for KO groups (Supplementary Table 8) and pathways (Supplementary Table 9). According to MaAsLin2, 50% of KO groups and 67% of pathways were affected in PD. According to ANCOM-BC, 32% of KO groups and 55% of pathways were affected. The overlap (i.e., FDR<0.05 by both methods) was 15% of KO groups and 32% of pathways. We therefore estimate that between 1/3 to 2/3 of all detected metabolic pathways are dysregulated in PD.

### Functional inference

Much of the constituents of the gut microbiome have been detected by metagenomics and not yet been characterized, nor can we infer on KO groups that have not yet been annotated. Thus, inferences made here are robust given the current state of knowledge and available resources, but do not capture the full potential of the study. While many of the pathways and KO groups that were altered in PD are non-specific and likely reflect broad dysbiosis, we noted several features in the PD metagenome that align with pathological features of PD (Fig. 3). Most functional insights to PD pathogenesis are derived from model organisms and in experimental setting, with the caveat that their relevance to human PD is not guaranteed. Here, we provide data from the human PD gut metagenome that corroborate, and validate, some of the basic science findings.

**Fig. 3.**
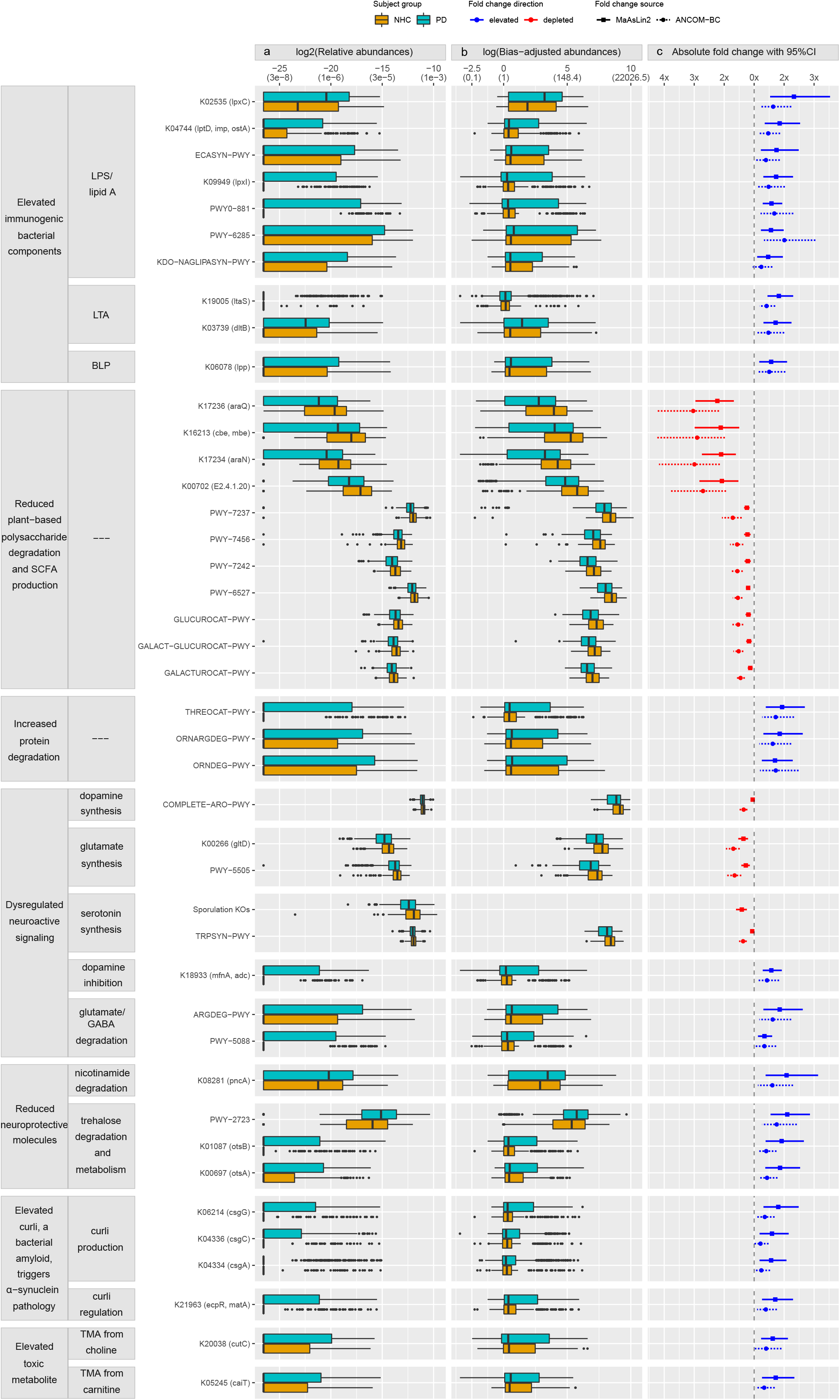
Microbial gene-families and pathways with functional relevance to PD. Overall, 15% of microbial gene-families (KO groups) and 30% of microbial pathways (MetaCyc) tested were elevated or depleted in PD, a conservative estimate derived from consensus at FDR<0.05 by two statistical methods (MaAsLin2 and ANCOM-BC) (Supplementary Tables 8, 9). Examples are shown here, grouped by inferred functional relevance to PD (left panel). Data show increased levels of microbial activities that could contribute to PD pathogenesis (immunogenicity, alpha-synuclein aggregation, and creation of toxic metabolites), and reduced levels of protective mechanisms (anti-inflammation, and depletion of neuroactive and neuroprotective molecules. The 3 columns of data correspond to **(a)** relative abundances (used in MaAsLin2), **(b)** bias-corrected observed abundances (used in ANCOM-BC), and **(c)** fold change in PD vs. NHC as estimated by MaAsLin2 and ANCOM-BC. <u>Y-axis:</u> KO groups (identifiers begin with “K”, gene symbol in parentheses) and pathways (“PWY”). <u>X-axis:</u> **(a)** Log2 transformed relative abundances, as used in MaAsLin2, **(b)** natural log transformed, sampling bias-corrected observed abundances, as used in ANCOM-BC, with untransformed relative abundances in parenthesis for easier interpretation; **(c)** Absolute fold change in differential abundance in PD vs. NHC with 95% confidence interval (CI), calculated from beta and standard errors estimated by MaAsLin2 and ANCOM-BC. <u>Content in (</u>**<u>a</u>** <u>and</u> **<u>b</u>**): Boxplots show frequency distribution of each KO and pathway in PD (blue green) and NHC (orange) metagenomes. Left, middle, and right vertical boundaries of each box represent first, second (median), and third quartiles of the data; that is, 25% of samples have abundance lower than the left border of the box, 25% of samples have abundances that are higher than the right border of the box. Absence of a box indicates >75% of samples had zero abundance. The lines extending from the two ends of each box represent 1.5x outside the interquartile range (range = (abundance value at 75% minus abundance value at 25%) x 1.5). Points beyond the lines are outlier samples. <u>Content in</u> **<u>(c):</u>** Fold change difference between PD and NHC in log2-transformed relative abundance (square with solid line of 95% CI, MaAsLin2), or natural log transformed bias-corrected observed abundances (circle with dotted line of 95% CI, ANCOM-BC). Blue: KO/pathway is elevated in PD, red: reduced in PD.

#### 1. Enrichment of Gram-negative bacteria, LPS, LPP, LTA gene-families and pathways

Inflammation has emerged as a major driver of PD pathogenesis^31^. We observed a general increase in PD metagenome of pro-inflammatory species, as well as gene-families and pathways that directly promote inflammatory signaling between microbes and the host. Lipopolysaccharides (LPS) are the most abundant antigen on the surface of Gram-negative bacteria, capable of eliciting strong immune reaction and inflammation, by stimulating the host’s Toll-like Receptor 4 (TLR4) signaling pathway. We detected a bloom of Gram-negative bacteria with immune-stimulatory LPS in PD metagenome. Eleven of the 55 species that are enriched in PD are canonical Gram-negative organisms including those with highly stimulatory LPS (*Escherichia coli, Klebsiella* species, and *Porphyromonas asaccharolytica*). Conversely, of the 29 species that are reduced in PD, only 1 is a canonical Gram-negative organism, *Prevotella copri*, which interestingly, produces an LPS that, not only does not induce inflammation, but can inhibit TLR4 activation by others^32^. We also detected elevated levels of gene-families and pathways necessary for generation and secretion of Lipid A, a glycolipid component of LPS responsible for much of its immune-stimulatory capacity^33^. The PD metagenome was enriched for the KO groups in the canonical (Raetz Pathway) lipid A synthesis pathway including *lpxC* (K02535: FC=2.3, FDR=1E-3), *lpxI* (K09949: FC=1.7, FDR=1E-3), *lptD* (K04744: FC=1.9, FDR=1E-3), and the lipid A synthesis pathway as a whole (KDO-NAGLIPASYN-PWY: FC=1.5, FDR=0.02). We further noted an enrichment of the synthesis pathway for the core, immunogenic sugar components of LPS (ECASYN-PWY: FC=1.7, FDR=5E-3), and the general fatty acid synthesis pathways necessary for production of the Gram-negative cell envelope (PWY-6285 and PWY0-881: FC=1.6, FDR=1E-3). We observed increases in gene-families involved in the synthesis of Lipoteichoic acid (LTA), a component of Gram-positive bacteria cell envelope which shares many of the pathogenic properties of LPS^34^ (K19005: FC=1.8, FDR=4E-5), and in its modification to be recognized by host TLR2 and stimulate inflammatory responses^35^ (K03739: FC=1.7, FDR=1E-3). Additionally, we observed enrichment in PD of murein/bacterial lipoprotein (BLP) gene-family [*lpp*] K06078 (FC=1.6, FDR=0.01). BLP is highly immune stimulatory by triggering inflammatory TLR2 signaling^36,37^, but is often overlooked comparatively to LPS.

#### 2. Reduction in species, genes and pathways that degrade polysaccharides and produce SCFA

SCFAs are products of fermentation of dietary fiber by the gut microbiome. SCFA affect host physiology through neuro-immune-endocrine signaling and epigenetic regulation of host genome^38^, and may play a key role in gut-brain communication^39^. In the gut, inadequate SCFA levels have been linked to constipation, compromised gut barrier, and inflammation^40,41^, all of which are common features of PD. This dataset revealed depletion of species, genes and pathways that metabolize plant-based polysaccharides and make SCFA. We observed a dramatic ∼2.5-fold reduction for several SCFA producing species, and as much as 5-fold reduction in *Roseburia* species. At gene-family level, many of the most decreased KO groups in the PD metagenome are those involved in polysaccharide utilization, each showing more than 2-fold reduction, e.g., *araQ* (K17236: FC=0.44, FDR=7E-6), *araN* (K17234: FC=0.48, FDR=9E-6), *cbe*/*mbe* (K16213: FC=0.47, FDR=4E-4), and E2.4.1.20 (K00702: FC=0.48, FDR=1E-4). We found a less dramatic, but statistically significant reduction in pathways critical for degradation and metabolism of plant-based polysaccharides including beta-mannan (PWY-7456: FC=0.81, FDR=1E-4), stachyose (PWY-6527: FC=0.83, FDR=3E-7), inositol (PWY-7237: FC=0.81, FDR=7E-7), fructuronate (PWY-7242 FC=0.83, FDR=1E-4), galacturonate (GALACTUROCAT-PWY: FC=0.89, FDR=3E-3), glucoronate (GLUCUROCAT-PWY: FC=0.84, FDR=1E-4), and galacturonate/glucoronate (GALACT-GLUCUROCAT-PWY FC=0.85, FDR=5E-4). Interestingly, the PD metagenome was enriched in glycogen metabolism pathway (GLYCOCAT-PWY: FC=2.0, FDR=7E-5), suggesting an enrichment in bacteria with a preference for non-plant-based polysaccharides. Of note, unlike most polysaccharide degradation pathways that are reduced, the starch degradation pathway III is enriched in the PD metagenome (PWY-6731: FC=2.0, FDR=7E-5), however, pathway III maps to *Escherichia coli* and *Klebsiella* species and is not known to produce beneficial SCFA.

#### 3. Enrichment in proteolytic pathways

Proteolytic and amino acid degradation pathways were substantially enriched in PD metagenome, suggesting PD microbiome preferentially utilizes proteins, over polysaccharides, as their prime carbon/energy source. All proteolytic pathways that were differentially abundant in PD vs. NHC were enriched in the PD metagenome, including pathways which degrade glutamate (PWY-5088: FC=1.34, FDR=3E-3), arginine/ornithine (ORNARGDEG-PWY: FC=1.85, FDR=2E-3, ARGDEG-PWY: FC=1.85, FDR=8E-3, ORNDEG-PWY FC=1.70, FDR=2E-3), and threonine (THREOCAT-PWY: FC=1.93, FDR=5E-4). These bacterial pathways, and specifically the upregulated threonine pathway, utilize protein sources derived from diet or the host, including mucin. Mucin is an important component of the protective gut mucus layer. Increased capacity to degrade host mucin raises the possibility that the gut microbiome contributes to the degradation and increased permeability of the gut barrier in PD^42^.

#### 4. Neuroactive molecules

Present data suggests substantial dysregulation in PD microbiome of synthesis and metabolism of small molecules that can modulate neuronal activity, and which are integral to PD pathologies, namely, dopamine, glutamate, gamma aminobutyric acid (GABA), and serotonin. Aberrations involving these neurotransmitters in PD is well-established^43^. The human gut microbiome has known capacity to not only produce and metabolize these neurotransmitters in the gut^44^, but also to influence their levels in the brain^45^.

##### Dopamine

A substantial drop in dopamine levels due to progressive loss of dopaminergic cells in the brain is the defining feature of PD. We detected elevated levels of tyrosine decarboxylase/aspartate 1-decarboxylase gene-family (K18933: FC=1.57, FDR=3E-4), which removes tyrosine, a required precursor of dopamine, thus limits dopamine production by host and bacteria. The aromatic amino acid synthesis pathway (COMPLETE-ARO-PWY: MaAsLin2 FC=0.95, FDR=6E-3; ANCOM-BC FC=0.74, FDR=1E-8) was depleted in PD metagenome, suggesting that the microbiome has diminished capacity to generate precursors to dopamine and possibly other catecholamines such as adrenaline.

##### GABA & Glutamate

The link between PD and glutamate toxicity due to overactivation of glutamate receptor has been demonstrated experimentally^46^ and associated to glutamate receptor gene in humans^47^. Imbalance between GABA and glutamate (excitation/inhibition imbalance) weakens synaptic connection and promotes neurodegeneration^48^. In the PD metagenome, we noted a decrease in glutamate/glutamine synthesis gene-families and pathways (K00266: FC=0.73, FDR=5E-5, PWY-5505: FC=0.78, FDR=1E-4), and a concomitant increase in both glutamate and GABA degradation (PWY-5088: FC=1.34, FDR=2E-3, ARGDEG-PWY FC=1.85, FDR=2E-3).

##### Serotonin

The highest concentration of serotonin is in the gut. Serotonin abnormalities have been detected at various stages of PD, from before the development of motor symptoms, making it a potential early diagnostic marker^49^, to late stages of PD where it may be a risk factor for cognitive decline^50^. It was shown in mice that spore forming bacteria induce production of serotonin in host colonic cells, which stimulates myenteric neurons and gut motility^51^. Tryptophan, an essential amino acid only derived from microbiome or diet, is the rate limiting precursor to serotonin. Tryptophan biosynthesis pathway was reduced in PD (TRPSYN-PWY: MaAsLin2 FC=0.94, FDR=9E-3; ANCOM-BC FC=0.73, FDR=7E-10). We also detected a near global reduction in sporulation genes in PD. We tested 33 sporulation KO groups in this dataset, 31 of them were depleted in PD (0.49 ≤ FC ≤ 0.77, 2E-5 ≤ FDR ≤ 0.04), suggesting that serotonin production in the host is depleted.

Given that spore forming bacteria stimulate gut motility^51^, and our data suggest a global depletion of the sporulation KO groups, we speculated, and tested the hypothesis that constipation, a common symptom of PD, may be related to the depletion of spore forming bacteria. We selected 27 sporulation KO groups that were FDR<0.05 in both ANCOM-BC and MaAsLin2 and tested the hypothesis that the relative abundance of sporulation KO groups is even lower in subjects who reported constipation vs. those that did not. Indeed, lower abundances of sporulation KO groups was associated with constipation both in PD group (FC=0.76, P=2E-4), and in NHC group (FC=0.58, FDR=3E-3). We should note that association of sporulation KO groups with PD is not solely due to constipation; adjusting for constipation, the relative abundance of sporulation KO groups was still significantly lower in PD than in NHC (FC=0.79, P=5E-4).

#### 5. Neuroprotective molecules

##### Nicotinamide

In PD, disruption of energy homeostasis leads to dysregulation of nicotinamide production and promotes neurodegeneration^52^. A recent clinical trial of PD with nicotinamide supplementation reported a significant improvement in pathological indicators including inflammation^53^. We observed a two-fold increase in the nicotinamidase gene-family (K08281: FC=2.1, FDR=3E-3), suggesting the PD gut microbiome degrades this neuroprotective molecule.

##### Trehalose

The disaccharide trehalose has garnered significant interest as a therapeutic molecule in PD and other neurodegenerative diseases^54^. This is likely due to its activities on autophagocytic pathways which limit the accumulation of pathogenic alpha-synuclein and other protein aggregates^55^. As we note with nicotinamide degradation, a similar two-fold increase was observed in the PD metagenome of the trehalose degradation pathway (PWY-2723: FC=2.1, FDR=4E-5). We also noted evidence for increased metabolism of trehalose by bacteria (K00697: FC=1.9, FDR=1E-3 and K01087: FC=1.9, FDR=1E-3), a potential outcome of which is reduced availability of trehalose for the host.

#### 6. Bacterial amyloid, curli

The bacterial-derived amyloidogenic protein, curli, exacerbates alpha-synuclein pathology and inflammation in experimental models of PD^17-19^. Many *Enterobacteriaceae* species encode curli^56^. We detected 33 species in *Enterobacteriaceae* family but only *Escherichia coli* and *Klebsiella* species had high-enough abundance to be tested, and they were all elevated in PD. We also found an enrichment of the gene-families related to curli. *csgA* (K04334: FC=1.6, FDR=9E-3), *csgB* (K04335: FC=1.3, FDR=0.04), *and csgC* (K04336: FC=1.6, FDR=0.01) are central to curli structure. *csgG* (K06214: FC=1.8, FDR=3E-3) is necessary for the extracellular secretion and synthesis of curli. LuxR/Mat/Ecp (K21963: FC=1.7, FDR=3E-3) is a family of transcriptional factors that regulate the expression of curli.

#### 7. Toxic metabolite

In humans, the gut microbiome produces trimethylamine (TMA) from foods such as red meat and egg yolk. High level of circulating TMA has been linked to increased risk of cardiovascular disease and stroke, likely due to toxic and inflammatory effects^57,58^. Circulating levels of TMAO, a gut microbiome derived oxidation product of TMA, are reportedly elevated in PD and correlate with disease progression and severity^59^. Gene-families involved in TMA production were elevated in PD, including *cutC* (choline lyase) which cleaves choline to produce TMA (K20038: FC=1.62, FDR=4E-3), and *caiT* (L-carnitine/gamma-butyrobetaine antiporter) which exchanges carnitine for gamma-butyrobetaine before TMA can be produced (K05245: FC=1.72, FDR=4E-3).

In summary, PD metagenome is indicative of a disease promoting microbiome (enriched in opportunistic pathogens and immunogenic components, dysregulated neuroactive signaling, preponderance of amyloidogenic molecules that induce alpha-synuclein pathology, and over production of toxicants), with reduced capacity for recovery (low in anti-inflammatory and neuroprotective molecules).

## Conclusion

We have generated, and share publicly, a large dataset at the highest resolution currently feasible. The dataset itself is a substantial contribution to open science because it includes individual-level deep shotgun metagenome sequences and extensive metadata on 490 persons with PD, the largest PD cohort with microbiome data, and a unique cohort of 234 neurologically healthy elderly, which can be used in a wide range of studies. We uncovered a widespread dysbiosis in PD metagenome that is indicative of an environment permissive for neurodegenerative events and prohibitive of recovery. This study exemplified bi-directional transfer of information between human and experimental studies: The data and results generated here from human metagenomes sets a broad foundation for basic research, including numerous concrete hypotheses which will be tested experimentally to discern the multitudes of roles that the gut microbiome may play in PD. At the same time, the results generated here confirmed, in humans, several observations that were made experimentally in animal models. While this study generated many hypotheses with solid evidence that can be tested now, it did not achieve the full potential of a metagenomics study. Metagenomics is a new, albeit fast evolving field, and the resources, methods, and tools, while state of the art, are still in development. Undoubtedly more information will be revealed as we increase the sample size, and others also conduct metagenomics studies and share the data. We anticipate that in near future we will have the tools and the analytic power to use metagenomics, as a new approach, to study PD heterogeneity, search for biomarkers, delve deeper into the origin and progression of PD sub-phenotypes (e.g., constipation, RBD), and investigate the potential in manipulating the microbiome to prevent, treat and halt the progression of PD.

## Methods

We have complied with all relevant ethical regulations. Study was approved by the Institutional Review Board (IRB) for Protection of Human subjects at the University of Alabama at Birmingham (UAB) and by the Human Research Protection Office (HRPO) of United States Department of Defense (funding agency). All subjects gave signed informed consent, approved by UAB IRB and DoD HRPO.

Study was conducted by a single team of NGRC investigators from start (study design and subject enrollment) to finish (analysis and interpretation), ensuring uniform methods. The same investigator team conducted the two prior studies that are being referenced here for comparison of shotgun and 16S analyses.

Software and reference databases used in this study are referred to by name only in the text for brevity; the versions and the URLs are provided in Supplementary Table 10. The workflow is summarized in Fig. 1.

### 1. Study cohort

#### 1.1. Enrollment

670 individuals with PD and 316 NHC were enrolled at UAB between October 2018 and March 2020. All subjects were from the same geographic region, including the city of Birmingham and surrounding areas in the southern US, keeping confounding effect of geography to a minimum. Potential PD cases for enrollment were identified via systematic pre-screening of electronic medical records (EMR) of patients with an upcoming appointment in the Movement Disorder Clinic at UAB. Subjects were invited to enroll in the study after their clinic visit, if the attending specialist confirmed PD diagnosis and the patient was willing to hear about the study. The eligibility criteria to enroll as a case was diagnosis of PD and informed consent. The spouse or friend accompanying the patient was invited to enroll as control, to match for shared environmental effects. Additional control subjects were recruited from the community. The eligibility criteria to enroll as a control was absence of PD, RBD, Alzheimer’s disease, dementia, multiple sclerosis, amyotrophic lateral sclerosis, ataxia, dystonia, autism, epilepsy, stroke, bipolar disorder, and schizophrenia by self-report (hence, neurologically healthy controls), preferably being over age 50 years, and informed consent. We tried not to enroll subjects who had participated in our prior microbiome studies to maintain cohort independence; however, 11 subjects were inadvertently double-enrolled. During enrollment visit, subjects signed an informed consent, donated a blood or saliva sample, and took home two questionnaires and a stool collection kit to complete and return in the pre-stamped envelope via US Postal Service. We ended the enrollment at the start of the COVID-19 pandemic in the southern US (March 2020) to avoid confounding by infection or stress of the pandemic.

#### 1.2. Exclusions

Subjects who did not return a stool sample were excluded (171 PD, 73 controls). One PD subject was excluded because the stool sample was received during the pandemic. We excluded 1 control for potential sample mix-up and 8 for having neurologic disease (7 stroke, 1 epilepsy). Since the diagnosis of PD can change with time, we reviewed EMR prior to sequencing, and again before data analysis, and those whose diagnosis changed were excluded (N=6). One PD subject was excluded for low sequence count. The final sample size was 490 PD and 234 NHC.

### 2. Metadata and stool sample collection

Subjects were given a packet containing two questionnaires (metadata), and a stool collection kit with detailed instructions and a self-addressed pre-stamped envelope for return.

#### 2.1. Metadata

The Environmental and Family History Questionnaire (EFQ) was used to collect PD-related data, and the Gut Microbiome Questionnaire (GMQ)^20,28^ was filled out by subject immediately after stool sample collection and gathered gut-related data. The questionnaires have been provided in Supplementary Fig. 1.

#### 2.2. Stool sample

Each subject provided one stool sample at one time point; hence each data point represents a unique individual. Each sample was measured and analyzed once. Stool samples were collected at home. Nearly all subjects (487 PD, 232 NHC) used the DNA Genotek (Ottawa, Ontario, Canada) OMNIgene GUT Collection kit. Five subjects (3 PD, 2 NHC) used Fisher Scientific (Hampton, NH, USA) DNA/RNA-free BD BBL Sterile/Media-free Swabs. Collection method was included as a technical covariate in analyses. Upon receipt in the lab, samples were stored in -20°C freezer.

#### 2.3. QC of returned packets

Upon receipt of each packet, the EFQ, GMQ and stool sample in the packet were cross-checked for potential labeling error in the lab or sample mix-up at home (PD and control pairs living together) and for completeness, comprehensibility, and lucidity of information. Issues were resolved by checking the EMR and calling the subject, and if not resolved, the sample or metadata in question were excluded. Metadata were entered in a customized PROGENY database software (Progeny Genetics LLC, Aliso Viejo, CA, USA), and in Excel spreadsheets, by two data entry staff, downloaded, cross-checked, and errors corrected. PROGENY was used for storage and retrieval of data.

### 3. DNA isolation, library preparation, and next-generation sequencing

Prior to shipping samples for processing, case and control samples were inter-mixed to avoid batch effect during DNA isolation and sequencing. Frozen stool samples were shipped on dry ice to the metagenomic services company CosmosID (Germantown, MD, USA). DNA was isolated from an aliquot of stool sample using the Qiagen (Germantown, MD, USA) DNeasy PowerSoil Pro high-throughput kit on the QIAcube high-throughput platform according to the manufacturer’s protocol. Isolated DNA was quantified by Qubit. The sequencing libraries were prepared by using the Illumina Nextera XT transposase system. Libraries were sequenced on the Illumina NovaSeq 6000 instrument aiming for 40M reads per sample, following paired-end 150bp sequencing protocol. The number of raw sequence reads per sample ranged within 17M to 385M (average 50M per sample).

### 4. Bioinformatic processing of sequences

#### 4.1. QC of sequences

The quality of sequencing was checked by FastQC, and aggregated quality reports for samples were generated using MultiQC^60^. QC of sequence reads was performed using BBDuk and BBSplit. In the first step of QC, BBDuk was used to remove Nextera XT adapter and PhiX genome contamination, and quality trim and filter sequences. BBDuk was ran with the ‘ftm=5’ to ensure any extra bases after the 150bp were removed, ‘tbo’ option to additionally trim adapters based on paired read overlap detection, ‘tpe’ option to trim paired reads to the same length, ‘qtrim=rl’ to quality trim both 5’ and 3’ ends of sequences, ‘trimq=25’ to trim ends of sequences up to quality score of 25, and ‘minlen=50’ to remove sequences that fall below 50bp after contaminate removal and trimming. Percent of sequences removed ranged from 22 to 72% of initial reads with a mean of 28% removed per sample. Next, all sequence reads mapping to human reference genome (GRCh38.p13) were removed using BBSplit with default parameters, eliminating <3% of BBDuk processed reads for most samples. Finally, we removed low-complexity sequences (<2% of human decontaminated reads) with BBDuk entropy filtering specifying ‘entropy=0.01’, which was performed to remove sequencing artifacts consisting of overrepresented long N-mers of “G”. The resulting QC’ed sequences consisted of 3M to 258M reads (average 36M) per sample. One sample with 3M reads was removed for analysis, but is included in the publicly available dataset. In essence, we followed the same QC protocol as recommended by HMP^61^, with one difference: as our sequencing protocol did not include PCR amplification step, we did not remove duplicated sequences, as these may be actual biological overrepresentation.

#### 4.2. Taxonomic profiling

QC’ed sequences were profiled using MetaPhlAn3^62^ with the accompanying database of ∼1.1M unique clade-specific marker genes. MetaPhlAn3 was ran twice: (1) relative abundances were generated using the default settings to be used in downstream analyses with MaAsLin2^63^, and (2) relative abundances were generated using the ‘--unknown-estimation’ flag to be multiplied by the total sequence reads of the sample to generate counts used in ANCOM-BC^64^. Taxonomic profiling resulted in 2,270 species represented by at least one marker gene, which were then trimmed by default parameters of MetaPhlAn down to 719 species.

#### 4.3. Enterotype profiling

Using relative abundances of genera, the web-based EMBL enterotype classifier was used to assign enterotypes to each sample based on the original enterotype definition (*Bacteroides, Firmicutes, Prevotella*)^65^. The frequency of each enterotype in PD and NHC were calculated separately, as number of samples assigned to each enterotype divided by total number of samples.

#### 4.4. Functional profiling

QC’ed sequences were profiled for potential functional content (UniRef90 gene-families^66^ and MetaCyc metabolic pathways^67^) using HUMAnN3^62^ with default parameters. Resulting UniRef90 gene-families were converted into KO groups using the HUMAnN utility tool ‘humann_regroup_table’. Pathways and KO groups were filtered to contain only community-level data. Functional profiling resulted in 8,528 distinct KO groups and 511 metabolic pathways, numbers which are in line with previous human gut microbiome studies^62^.

### 5. Statistical analysis

The sample size for all statistical analyses included 490 PD and 234 NHC, except confounding analysis, which included 487 PD and 232 NHC. When using R, default parameters were used unless noted. Every test, except SparCC correlations, included technical variables as covariates. Technical variables were (1) total sequence count per sample (standardized using ‘scale’ in R), and (2) stool collection kit (5 subjects used sterile swabs, 719 used OMNIgene GUT). Other covariates were added as appropriate and will be noted. All P-values were two-sided.

#### 5.1. Analysis of metadata

Distributions of 53 variables (Table 1) in PD vs. NHC were tested using Fisher’s exact test for categorical variables with ‘fisher.test’ in R, and using Wilcoxon rank-sum test for quantitative variables with ‘wilcox.test’ in R. ORs and 95% confidence intervals (CI) were calculated using ‘fisher.test’ unless the 2×2 table contained a zero, then ‘Prop.or’ from the pairwiseCI R package specifying ‘CImethod=‘Woolf’’ was used. P-values were not corrected for multiple testing because the aim was to detect any trend of a difference that could point to a potential confounder.

#### 5.2. Testing the overall composition of the metagenome

##### 5.2.1. Principal component analysis (PCA)

We used Aitchison distances as the measure of inter-sample differences in the compositions of gut metagenomes. Aitchison distances are Euclidean distances calculated from centered log-ratio (clr) transformed species count data. The clr transformation was performed using equation (1) in R: clr transformed species counts = log(*x*+1) - mean(log(*x*+1)) where *x* is a vector of all species counts in a sample. PCA was performed on Aitchison distances, using the ‘prcomp’ in R. The first two PC’s were plotted using ‘autoplot’ from the ggfortify R package.

##### 5.2.2. Beta-diversity

The difference in the overall composition of PD vs. NHC metagenomes was formally tested on Aitchison distances using PERMANOVA^68^ via ‘adonis2’ from the vegan R package. Significance of PERMANOVA was determined using 9,999 permutations.

##### 5.2.3. Enterotypes

Differences in the enterotype frequencies in PD vs NHC was tested using Chi-squared test via ‘chisq.test’ in R. Frequency distribution of enterotypes in PD and NHC were plotted using ‘mosaic’ from the vcd R package.

#### 5.3. Differential abundance analysis

##### 5.3.1. Unbiased differential abundance analysis: MWAS

MWAS approach was used to test differential abundances of species, genera, KO groups, and pathways in turn, in 490 PD vs. 234 NHC. Each test included all detected features that passed sequence QC and which were present in >5% of subjects (257 species, 107 genera, 4,679 KO groups, 407 pathways). The primary analysis was species-level MWAS. Genus-level MWAS was for comparability to past literature. KO group and pathway MWAS were for functional inference.

- *Statistical tests*. Two multivariable association tests, MaAsLin2 and ANCOM-BC, were used in parallel to ensure results were robust to methodological variations. We chose MaAsLin2 and ANCOM-BC because in comparative studies with real and simulated data they were among the most robust in keeping false positive rate low while maintaining power^69,70^. MaAsLin2 is the differential abundance test in the bioBakery suite of tools, which we used for much of the bioinformatics. ANCOM-BC differs in approach, and in addition, has introduced the notion of “sampling fraction” to estimate how well the observed abundances in a sample represent the true absolute abundances in the ecosystem (human gut) and incorporates bias correction to adjust for differences in sampling fractions across samples. Thus, since MaAsLin2 tests relative abundances, and ANCOM-BC tests bias-corrected observed abundances (approximating absolute abundances), the results shown side-by-side provide different and complementary perspective on how PD and NHC metagenomes differ.
- *Multiple testing correction*. Benjamini-Hochberg FDR method^71^ was used.
- *Significance*. For species-level MWAS (the primary analysis), we set statistical significance for association with PD as concordance between ANCOM-BC and MaAsLin2 with FDR<0.05 by one and FDR≤0.1 by the other. For other MWAS, while no hard rule was set, results discussed are those that are in the FDR ∼0.05 to ∼0.1 range.
- *Effect sizes*. Measured as fold change (FC) and calculated in R using equation (2) and (3) for ANCOM-BC and MaAsLin2 respectively:

(2) FC = exp(*x*)

(3) FC = 2^ (*x*)

where *x* is a vector of model coefficients derived from ANCOM-BC (based on natural log transformed abundances), or MaAsLin2 (based on log2-transformed relative abundances).

- *Plots*. Relative abundances (MaAsLin2) and bias-corrected observed abundances (ANCOM-BC) in PD and NHC, and the FCs with 95% CI were plotted for PD-associated species and selected KO groups and pathways using ggplot2 R package. To extract bias-corrected observed abundances from ANCOM-BC to plot, we used equation (4) in R: (4) Bias-corrected observed abundances = exp(log(*x* + 1) – *y*) where *x* is a sample by feature matrix of observed abundances, and *y* is a vector of sampling fractions estimated by ANCOM-BC for each sample.
- *Parameters*. MaAsLin2 and ANCOM-BC were run with default parameters, except: microbial features that were not present in ≥5% of samples were excluded from testing to ensure minimum analytic N=37 (‘zero_cut=0.95’ for ANCOM-BC, ‘min_prevalence=0.05’ for MaAsLin2), and multiple testing correction was Benjamini-Hochberg FDR^71^ (‘p_adj_method=“BH”’ for ANCOM-BC, ‘correction=“BH”’ for MaAsLin2). Additionally, for MaAsLin2, ‘normalization’ was set to “NONE” as data input was already total sum scaled normalized, and ‘standardize’ was set to “FALSE” as standardization of quantitative variables was done prior to running MaAsLin2.
- *Confounder analysis*. To investigate the confounding effects of extrinsic variables that differed between PD and NHC, MaAsLin2 was performed with the 84 PD-associated species from MWAS, and 9 variables in the model: PD/NHC, total sequence count, intake (yes/no) of alcohol, laxatives, probiotics, antihistamines, depression-anxiety-mood medication, pain medication, and sleep-aid. Five subjects were excluded because they were missing pain medication and sleep-aid data. Stool collection method was not included as the 5 excluded subjects were those who used swabs. Although PD and NHC also differed in weight loss, GI discomfort, and constipation, these variables were not considered confounders because they are intrinsic features of PD. Adjusting for them would mask a part of PD, which is counter to the aim of this study: to generate a full unaltered view of the dysbiosis in PD metagenome. Age and sex are also intrinsic to PD as they are risk factors. However, the selection of household controls to minimize environmental variation inflated the female preponderance and younger ages in controls as PD is more prevalent in men. To ensure the species-level GWAS results were not artifacts of this distortion, we ran MaAsLin2 on the 84 PD-associated species from MWAS and 5 variables in the model: PD/NHC, total sequence count, stool collection method, male/female, and age at stool collection (in years and standardized using ‘scale’ in R).

##### 5.3.2. Hypothesis-driven differential abundance analyses

The following tests were conducted using linear regression (via ‘lm’ in R) on log2-transformed relative abundances as done with MaAsLin2.

- *Prevotella sub-genus*. This test was conducted to assess reproducibility of an earlier finding of elevated abundances of a *Prevotella* sub-genus^20^ that was identified by the 16S database SILVA, and included *Prevotella buccalis, timonensis, bivia, disiens* and *oralis*. At species-level they were individually too rare and excluded from MWAS. We collapsed relative abundances of these species to create the *Prevotella* sub-genus and tested the difference in PD vs. NHC.
- *Cluster #17*. This test was conducted to assess reproducibility of an earlier finding of elevated abundances of opportunistic pathogens detected at genus-level^20^. Again, individual species were rare and not tested, except *Porphyromonas asaccharolytica*. We combined them, as was done in prior study with genera, and tested the difference between PD and NHC. Test was performed once with all 19 species of cluster #17 combined (listed in Supplementary Table 7), and once excluding *Porphyromonas asaccharolytica*.
- *Sporulation KO groups and constipation*. This test was conducted because sporulation KO groups were reduced in PD, and prior studies have shown sporulation species are necessary for gut motility^51^; hence the question: Does our data support an association between reduced levels of sporulation KO groups and constipation? To test this, we collapsed the 27 sporulation KO groups that had shown association with PD (i.e., in Supplementary Table 8, searched KO group names for key word “sporulation”, and selected those with FDR<0.05 by ANCOM-BC and MaAsLin2), and tested their combined relative abundance in constipated (in past 3 months) vs. non-constipated strata, within PD and NHC separately. To determine if sporulation KO groups were significantly depleted in PD independently of constipation, we re-tested the association of case status with abundance of sporulation KO groups, this time adjusting for constipation in the model.

#### 5.4. Network analysis

Correlation networks were generated for PD and NHC microbiomes separately. Pairwise SparCC correlations^72^ were calculated using species count data as input to ‘fastspar’ from FastSpar (a C++ implementation of SparCC, ^73^) specifying 100 iterations. Permuted P-values were then calculated for each correlation by (1) creating 1,000 random datasets, (2) calculating SparCC correlations for each random dataset, and (3) calculating P-values for each correlation by determining the proportion of random correlations that were stronger than the original correlations. The Louvain algorithm for community detection^74^ was used (via ‘cluster_louvain’ from the igraph R package) to algorithmically detect clusters of species using correlations that reached |r| > 0.2 and permuted P < 0.05. The number of correlations at |r| > 0.2 and permuted P < 0.05 for each species was calculated using ‘degree’ from igraph. To visualize networks, correlations that reached |r| > 0.2 with permuted P<0.05 were imported into Gephi along with their cluster memberships. The Force Atlas 2 algorithm^75^ was then used to plot the network, showing species as nodes, correlations as connecting edges between species, and strength of correlation by the weight of the edge.

## Data availability

Individual level raw shotgun metagenomic sequences and metadata are publicly available at NCBI Sequence Read Archive (SRA) under BioProject ID PRJNA834801 (*will be released upon publication*).

## Code availability

Code used to perform bioinformatics can be found at GitHub (https://github.com/zwallen/SLURM_Shotgun_Metagenomic_Pipeline) and at Zenodo with DOI: https://doi.org/10.5281/zenodo.6588381. Code and input data used to perform the statistical analyses can be found at GitHub (https://github.com/zwallen/Wallen_et_al_PDShotgunAnalysis) and at Zenodo with DOI: https://doi.org/10.5281/zenodo.6620579.

## Acknowledgement

Authors thank individuals who participated in this research project, and the movement disorder specialists at UAB. This work was supported by The U.S. Army Medical Research Materiel Command endorsed by the U.S. Army through the Parkinson’s Research Program Investigator-Initiated Research Award under Award No. W81XWH1810508 (to HP); NIH Training Grant T32NS095775 (to ZDW); NIH/NIEHS 1R01ES032440-01A1 and Parkinson’s Foundation PF-SF-JFA-830658 (to TS); and Aligning Science Across Parkinson’s [ASAP-020527] through the Michael J. Fox Foundation for Parkinson’s Research (MJFF) (to HP and TS). For the purpose of open access, the authors have applied a CC BY public copyright license to all Author Accepted Manuscripts arising from this submission. Opinions, interpretations, conclusions, and recommendations are those of the authors and are not necessarily endorsed by the funding agencies.

## Author contribution

All authors met all four criteria (1) substantial contributions to the conception (H.P.) or design (H.P., Z.D.W.) of the work or the acquisition (H.P., D.G.S., M.N.D.), analysis (Z.D.W., G.C., G.T., A.D., H.P.) or interpretation of the data (H.P., T.S., Z.D.W.); (2) drafting the work (H.P., Z.D.W., T.S., A.D.) or revising it critically for important intellectual content (all authors); (3) final approval of the completed version (all authors); (4) accountability for all aspects of the work in ensuring that questions related to the accuracy or integrity of any part of the work are appropriately investigated and resolved (all authors).

