## Supplementary Figures for "Metagenomics of Parkinson’s disease implicates the gut microbiome in multiple disease mechanisms"

**Supplementary Fig. 1. Metadata questionnaires.** Following STORMs reporting guidelines for human microbiome research (PMID: 34789871), we are providing the questionnaires in full and have highlighted the questions whose data were used as metadata in the current study. Questionnaires and a stool sampling kit were given to the subjects to take home, self-complete, and return via postal service.

**For Research Laboratory Use Only**

Subject ID: ________________ PD____ RBD___ Control__ Day sample received M____ D_____ Y_____

*Interactions of Gut Microbiome, Genetic Susceptibility, and Environmental Factors in Parkinson's Disease*

A Research Study Funded by The United States Department of Defense

### GUT MICROBIOME QUESTIONNAIRE

Thank you for participating in this research study. Please complete this form immediately after collecting the stool sample and send it back with the stool sample and the completed Environmental & Family History Questionnaire in the enclosed pre-stamped envelope. If you have any questions, please call 205-934-0371.

Today's date (day of stool sample collection): Month__________ Day________ Year_________

Your Name: ________________________________________________________

Sex: ❒ M ❒ F

Birthdate: Month ______ Day________ Year ________

Phone _________________________________ Email _________________________________

On the **DAY OF STOOL COLLECTION** did you have:

Abdominal pain or discomfort ❒No ❒Yes

Bloating ❒No ❒Yes

Diarrhea ❒No ❒Yes

Excess gas ❒No ❒Yes

Constipation (no bowel movement for 3 days prior to stool collection) ❒No ❒Yes

In this **Bristol Stool Chart**, circle the type of stool that you passed today from which you collected the sample:

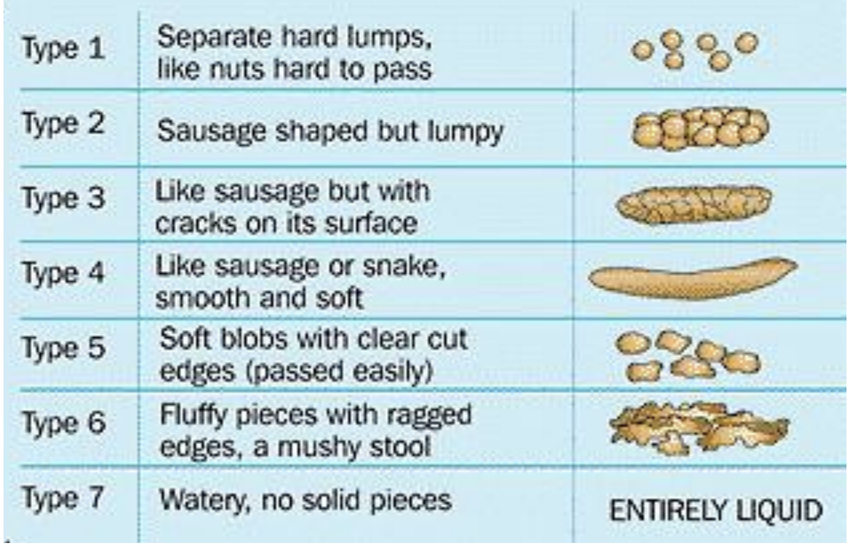

How tall are you? Feet_____ Inches_________

How much do you weigh? Pounds_________

Have you lost more than 10 pounds in the last year? ❒ No ❒ Yes

Have you gained more than 10 pounds in the last year? ❒ No ❒ Yes

In a typical week, how many hours of sleep do you get each night? ___________ hours

Do you smoke cigarettes, cigars or a pipe? ❒ No ❒ Yes

Do you drink caffeinated coffee, caffeinated tea, or caffeinated soda? ❒ No ❒ Yes

Do you drink alcohol? ❒ No ❒ Yes

#### DIET

How often do you eat **GRAINS** (rice, bread, pasta)?

❒ At least once a day
❒ Few times a week
❒ Few times a month
❒ Less than once a month or never

How often do you eat **POULTRY, BEEF, PORK, SEAFOOD, EGGS?**

❒ At least once a day
❒ Few times a week
❒ Few times a month
❒ Less than once a month or never

How often do you eat **FRUITS or VEGETABLES**?

❒ At least once a day
❒ Few times a week
❒ Few times a month
❒ Less than once a month or never

How often do you eat **NUTS**?

❒ At least once a day
❒ Few times a week
❒ Few times a month
❒ Less than once a month or never

How often do you eat **YOGURT**?

❒ At least once a day
❒ Few times a week
❒ Few times a month
❒ Less than once a month or never

#### MEDICAL CONDITIONS

Do you have Parkinson's disease? ❒No ❒Yes

Do you have rapid eye movement sleep behavior disorder (RBD)? ❒No ❒Yes

Are you often constipated (fewer than 3 bowel movements per week occurring frequently)? ❒No ❒Yes

Do have Diarrhea often (once a week or more, type 7 in Bristol stool chart)? ❒No ❒Yes

Do you have Irritable Bowel Syndrome (IBS) or spastic colon? ❒No ❒Yes

Do you have Inflammatory Bowel Disease (IBD)? ❒No ❒Yes

Do you have Small Intestinal Bacterial Overgrowth (SIBO)? ❒No ❒Yes

Have you had an Ulcer in the past three months? ❒No ❒Yes

Do you have Celiac disease? ❒No ❒Yes

Do you have Crohn's disease? ❒No ❒Yes

Do you have Colitis? ❒No ❒Yes

Have you had a cancer of the digestive system in the last 3 months? ❒No ❒Yes

#### MEDICATIONS

Are you currently taking antibiotics? ❒ No ❒ Yes

Have you completed a course of antibiotics in the past 3 months? ❒ No ❒ Yes

Do you take laxatives at least once a week? ❒ No ❒ Yes

Do you take drugs for indigestion or reflux at least once a week? ❒ No ❒ Yes

Do you take anti-inflammatory drugs at least once a week? ❒No ❒Yes

Are you currently being treated for cancer with radiation or chemotherapy? ❒ No ❒ Yes

Are you taking blood thinners? ❒ No ❒ Yes

Are you taking cholesterol medication? ❒No ❒Yes

Are you taking blood pressure medication? ❒ No ❒ Yes

Are you taking thyroid medication? ❒ No ❒ Yes

Are you taking medication for asthma or COPD? ❒ No ❒ Yes

Are you taking medication for diabetes, high blood sugar, insulin? ❒ No ❒ Yes

Are you taking pain medication? ❒ No ❒ Yes

Are you taking medication for depression, anxiety, mood? ❒ No ❒ Yes

Are you on birth control pills or estrogen replacement therapy? ❒ No ❒ Yes

Are you taking Antihistamines? ❒ No ❒ Yes

Do you take Probiotic supplements? ❒ No ❒ Yes

Do you take Co-Q 10 supplements? ❒ No ❒ Yes

Do you take a sleep aid to help you fall asleep or stay asleep? ❒ No ❒ Yes

Are you **CURRENTLY** taking any **PARKINSON MEDICATIONS?** ❒No ❒Yes

If no, you can skip the rest of the form.
If yes, check all medications that you are taking (we have given the two names available for each medication).

**Levodopa preparations:**

❒ No ❒ Yes Sinemet or immediate release Carbidopa-Levodopa mg/pill_____Number of Pills/day_____

❒ No ❒ Yes Sinemet CR or Carbidopa/Levodopa ER mg/pill_____Number of Pills/day_____

❒ No ❒ Yes Rytary or Carbidopa/levodopa Capsules mg/pill_____Number of Pills/day_____

❒ No ❒ Yes Duopa or levodopa intestinal gel mg/24 hrs_____

❒ No ❒ Yes Stalevo or Carbidopa/levodopa/entacapone mg/pill_____Number of Pills/day_____

**Dopamine agonist drugs:**

❒ No ❒ Yes Mirapex or Pramipexole immediate release or ER mg/pill_____Number of Pills/day_____

❒ No ❒ Yes Requip or Ropinirole immediate release or XR mg/pill_____Number of Pills/day_____

❒ No ❒ Yes Neupro or Rotigotine patch mg/patch_____

❒ No ❒ Yes Apokyn or apomorphine injections or infusion cc/injection_____Number of injections/day_____

**COMT Inhibitors:**

❒ No ❒ Yes Comtan or Entacapone mg/pill_____Number of Pills/day_____

❒ No ❒ Yes Tasmar or tolcapone mg/pill_____Number of Pills/day_____

**MAO Inhibitors:**

❒ No ❒ Yes Azilect or Rasagiline mg/pill_____Number of Pills/day_____

❒ No ❒ Yes Eldepryl or Deprenyl or Selegiline mg/pill_____Number of Pills/day_____

❒ No ❒ Yes Xadago or Safinamide mg/pill_____Number of Pills/day_____

**Anticholinergic drugs:**

❒ No ❒ Yes Artane or Trihexyphenidyl mg/pill_____Number of Pills/day_____

❒ No ❒ Yes Cogentin or Benztropine mg/pill_____Number of Pills/day_____

**Other medication for Parkinson:**

❒ No ❒ Yes Symmetrel or Amantadine mg/pill_____Number of Pills/day_____

**Other Parkinson medications:**

1. Medication name ______________________ mg/pill_____Number of Pills/day_____
2. Medication name______________________ mg/pill_____Number of Pills/day_____
3. Medication name______________________ mg/pill_____Number of Pills/day_____

**Thank you for completing the questionnaire.**
**Please mail it back with the stool sample and the Environmental & Family History Questionnaire using the pre-stamped envelope. You may drop the envelope at any US postal service box.**

*Interactions of Gut Microbiome, Genetic Susceptibility, and Environmental Factors in Parkinson's Disease*

A Research Study Funded by The United States Department of Defense

### ENVIRONMENTAL & FAMILY HISTORY QUESTIONNAIRE

Thank you for participating in this research study. Please complete this form and send it back with the stool sample and the completed Gut Microbiome Questionnaire in the enclosed pre-stamped envelope. If you have any questions, please call 205-934-0371.

Today's Date: ____________________

Name: __________________________

Sex: ❒ M ❒ F

Birthdate: Month _____ Day_______ Year _______

Phone: (______)________________________ Email: ______________________________

Address: ______________________________ City_____________ State________ Zip Code __________

#### MEDICAL CONDITIONS

diagnosis was by patient's movement disorder specialist

Do you have **Parkinson's disease**? ❒ No ❒ Yes
How old were you when you first noticed a sign of Parkinson's disease (age at onset)? ______years old
At what age did you receive the diagnosis of Parkinson's disease (age at diagnosis)? ______years old

Do you have **Rapid eye movement sleep Behavior Disorder (RBD)**? ❒ No ❒ Yes
At what age did you receive the diagnosis of RBD? ______years old

Have you had a stroke, ataxia, multiple sclerosis, Alzheimer's disease, dementia, dystonia, autism, bipolar disorder, amyotrophic lateral sclerosis (ALS), or epilepsy? ❒No ❒Yes

- If Yes, circle the disorder.

Exclusion criteria for controls

#### HEAD INJURY

Have you had a head injury that caused loss of consciousness or required medical care? ❒No ❒Yes
If yes, how old were you when it first happened? ______________ years old

Did you have repeated blows to the head such as in sports or military? ❒No ❒Yes

#### TOXINS

Were you ever exposed to Agent Orange or other chemical warfare? ❒No ❒Yes ❒Don't know

Were you ever exposed to heavy uses of pesticides or herbicides, for example, did you live on or near farms that did crop dusting? ❒No ❒Yes
If yes, how old were you (give the range, for example, from birth until age 16) ___________________

#### TOBACCO

Have you smoked at least 100 cigarettes (about 5 packs) in your entire lifetime? ❒Yes ❒No
If you never smoked at least 100 cigarettes, go to "CAFFEINE" section.

During the time that you smoked, how much did you smoke on average, and for how many years? Check all that apply.

❒Less than ½ pack per day, for _____ years *(specify number of years)*
❒Equal to or more than ½ pack but less than 1 pack per day, for _____ years
❒Equal to or more than 1 pack but less than 2 packs per day, for _____ years
❒Equal to or more than 2 packs per day, for _____ years

At what age did you begin smoking? _________

Are you still smoking? ❒Yes ❒No If not, at what age did you stop? _______________

How many cigarettes do you currently smoke?

❒None
❒Less than ½ pack per day
❒Equal to or more than ½ pack but less than 1 pack per day
❒Equal to or more than 1 pack but less than 2 packs per day
❒Equal to or more than 2 packs per day

#### CAFFEINE

How much <u>caffeinated **COFFEE**</u> do you (or did you) drink, and for how many years?
*A **cup** is the size of a china cup. A **mug** is 2 cups.* Check all that apply.

❒None
❒Less than 2 cups a week, for _____ years *(specify number of years)*
❒Equal to or more than 2 but not more than 6 cups a week, for _____ years
❒1-2 cups a day, for _____ years
❒3-5 cups a day, for _____ years
❒6 or more cups a day, for _____ years

At what age did you start drinking caffeinated coffee? _______ years old

Are you still drinking caffeinated coffee? ❒Yes ❒No If not, at what age did you stop? ________

How much caffeinated coffee do you currently drink?

❒None
❒Less than 2 cups a week
❒Equal to or more than 2 but not more than 6 cups a week
❒1-2 cups a day
❒3-5 cups a day
❒6 or more cups a day

How much <u>caffeinated **TEA**</u> do you (or did you) drink and for how many years? *(A **cup** is the size of a china cup. A **mug** is 2 cups).* Check all that apply.

❒None
❒Less than 2 cups a week, for _____ years *(specify number of years)*
❒Equal to or more than 2 but not more than 6 cups a week, for _____ years
❒1-2 cups a day, for _____ years
❒3-5 cups a day, for _____ years
❒6 or more cups a day, for _____ years

At what age did you start drinking caffeinated tea? _______

Are you still drinking caffeinated tea? ❒Yes ❒No If not, at what age did you stop? ________

How much caffeinated tea do you currently drink?

- ❒None
- ❒Less than 2 cups a week
- ❒Equal to or more than 2 but not more than 6 cups a week
- ❒1-2 cups a day
- ❒3-5 cups a day
- ❒6 or more cups a day

How much caffeinated **SODA** do you (or did you) drink and for how many years? Check all that apply.

- ❒None
- ❒Less than 2 cans a week, for _____ years *(specify number of years)*
- ❒Equal to or more than 2 but not more than 6 cans a week, for _____ years
- ❒1-2 cans a day, for _____ years
- ❒3-5 cans day, for _____ years
- ❒6 or more cans a day, for _____ years

At what age did you start drinking caffeinated soda? _______

Are you still drinking caffeinated soda? ❒Yes ❒No If not, at what age did you stop? ________

How much caffeinated soda do you currently drink?

- ❒None
- ❒Less than 2 cans a week
- ❒Equal to or more than 2 but not more than 6 cans a week
- ❒1-2 cans a day
- ❒3-5 cans a day
- ❒6 or more cans a day

#### ALCOHOL

How much alcohol do you (or did you) drink, and for how many years? 1 drink is a beer, or a glass of wine, or a shot of liquor. Check all that apply.

- ❒Never
- ❒Less than 2 drinks a week, for _____ years *(specify number of years)*
- ❒2-6 drinks a week, for _____ years *(specify number of years)*
- ❒1 drink a day, for _____ years *(specify number of years)*
- ❒2 drinks a day, for _____ years *(specify number of years)*
- ❒3 or more drinks a day, for _____ years *(specify number of years)*

At what age did you start drinking alcohol? _______

Are you still drinking? ❒Yes ❒No If not, at what age did you stop? ________

How much alcohol do you currently drink?

- ❒Not at all
- ❒Less than 2 drinks a week
- ❒2-6 drinks a week
- ❒1 drink a day
- ❒2 drinks a day
- ❒3 or more drinks a day

#### NSAIDs

NSAIDs are non-steroidal anti-inflammatory drugs, like Ibuprofen, Motrin IB, Advil and Aleve, which are commonly used for pain. Aspirin and acetaminophens like Tylenol are not NSAIDs.

How often do you (or did you) take over the counter NSAIDs (like Ibuprofen, Motrin IB, Advil and Aleve), and for how many years? Check all that apply

- ❒Never
- ❒Less than once a week, for _____ years *(specify number of years)*
- ❒About one to four times a week, for _____ years
- ❒About 5 to 10 times a week, for _____ years
- ❒More than 10 times a week, for _____ years

How old were you when you started taking over the counter NSAIDs?_______

How often do (or did) you take prescription NSAIDs, and for how many years? Check all that apply.

- ❒Never
- ❒Less than once a week, for _____ years *(specify number of years)*
- ❒About one to four times a week, for _____ years
- ❒About 5 to 10 times a week, for _____ years
- ❒More than 10 times a week, for _____ years

How old were you when you started taking prescription NSAIDs? _______

#### OCCUPATION

Please indicate the types of work you have done over your lifetime, Check all that apply and give the year you started and stopped each occupation. If you don't remember the exact years, give us your best estimate.

| | Year started | Year ended |
|---|---|---|
| ❒Agriculture | ______________ | ______________ |
| ❒Gas | ______________ | ______________ |
| ❒Electricity | ______________ | ______________ |
| ❒Water and sewer | ______________ | ______________ |
| ❒Transportation | ______________ | ______________ |
| ❒Mining | ______________ | ______________ |
| ❒Construction | ______________ | ______________ |
| ❒Manufacturing | ______________ | ______________ |
| ❒Physician or nurse | ______________ | ______________ |
| ❒Office job | ______________ | ______________ |
| ❒Other | | |
| ❒Did not work outside the house | | |

#### RESIDENCE

Please tell us about all the places you have lived from birth to your current home address.

Give the years you were at each place (If you don't remember the exact years, give your best estimate.)

| Year moved in | Year moved out | Town | State | Country |
|---|---|---|---|---|
| Birth year | ___________ | _______________ | ___________ | ___________ |
| __________ | ___________ | _______________ | ___________ | ___________ |
| __________ | ___________ | _______________ | ___________ | ___________ |
| __________ | ___________ | _______________ | ___________ | ___________ |
| __________ | ___________ | _______________ | ___________ | ___________ |
| __________ | ___________ | _______________ | ___________ | ___________ |
| __________ | ___________ | _______________ | ___________ | ___________ |
| __________ | ___________ | _______________ | ___________ | ___________ |

#### FAMILY HISTORY

From what country did your **father's ancestors** immigrate to the US? ________________

From what country did your **mother's ancestors** immigrate to the US? ________________

Are you Hispanic or Latino? ❒ No ❒Yes

What race do you most identify yourself with?

❒White ❒Black or African American ❒American Indian/Alaskan Native

❒Asian ❒Native Hawaiian or other Pacific Islander ❒More than one race

Are you of Jewish ancestry? ❒No ❒Yes

If you are from a particular ethnic or religious lineage please specify. This is a question of heritage and genetic lineage, not personal preference: ____________________

Do you have any blood relatives who have **Parkinson's disease** (including your parents, grand-parents, siblings, aunts and uncles)? ❒No ❒Yes

If yes, for each relative who has or had PD, list their relationship to you.

(1) ________________________ (2) ________________________

(3) ________________________ (4) ________________________

Do you have any blood relatives who have **rapid eye movement sleep behavior disorder (RBD)** (including your parents, grand-parents, siblings, aunts and uncles)? ❒No ❒Yes

If yes, for each relative who has or had RBD, list their relationship to you.

(1) ________________________ (2) ________________________

(3) ________________________ (4) ________________________

**Thank you for completing the questionnaire.**

**Please mail it back with the stool sample and the Gut Microbiome Questionnaire using the pre-stamped envelope. You may drop the envelope at any US postal service box.**

**Supplementary Fig. 2. Global composition of gut metagenome in PD differs from NHC.** **(a)** Principal component analysis using Aitchison distances between 490 PD and 234 NHC metagenomes. While the difference between PD and NHC was not visually discernible, formal statistical test of inter-sample differences (beta-diversity), measured as Aitchison distance and tested using PERMANOVA, showed they significantly differed (P < 1E-4; 9,999 permutations, adjusted for total sequence count per sample and stool sample collection kit). **(b)** Enterotype profiles for PD and NHC groups were also significantly different (P = 4E-4), driven by the enrichment of *Firmicutes* in PD, and to a lesser extent a reduction in *Prevotella*. The height of each row represents the sample size, and the width represents the percentage of samples classified as that enterotype.

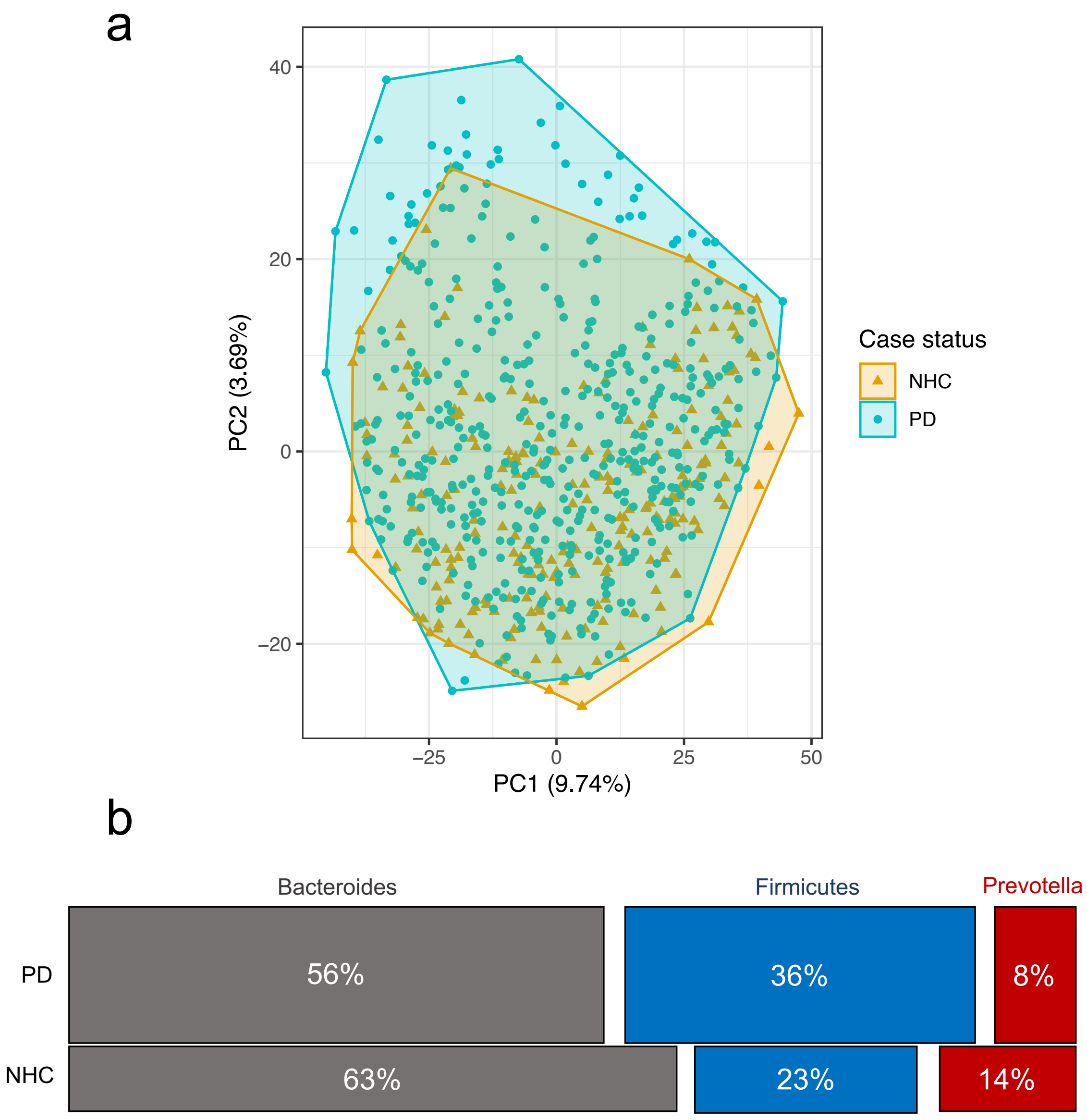

**Supplementary Fig. 3. PD-associated species' abundance distribution and fold change.** Relative abundances, bias-corrected observed abundances, and fold changes of 84 species that emerged from MWAS as having significantly elevated or significantly reduced abundances in PD vs. NHC were plotted. In MWAS, significance was defined as concordance between MaAsLin2 and ANCOM-BC with FDR<0.05 in one and FDR<0.1 in the other. **(a)** Distribution of relative abundances. Log2 transformed relative abundance values, as used in MaAsLin2, were used here to generate the boxplots. Untransformed relative abundances, shown in parenthesis, are provided on the x-axis for easier interpretation of data. Boxplots show distribution of the data for PD (blue green) and NHC (orange). Each sample was plotted according to its abundance of the species. The left, middle, and right vertical boundaries of each box represent the first, second (median), and third quartiles of the data; that is, 25% of samples have abundance lower than the left border of the box, 25% of samples have abundances that are higher than the right border of the box. Absence of a box indicates 75% of samples had zero abundance. The lines extending from the two ends of each box represent 1.5x outside the interquartile range (range = (abundance value at 75% minus abundance value at 25%) x 1.5). Points beyond the lines are outlier samples. **(b)** Distribution of bias-corrected observed abundances (used in ANCOM-BC). Natural log transformed, sampling-bias corrected observed abundances, as used in ANCOM-BC, were used here to generate the boxplots. Untransformed bias-corrected observed abundances, shown in parenthesis, are provided on the x-axis for easier interpretation of data. Description of boxplots are the same as for (a). **(c)** Absolute fold change in differential abundance in PD vs. NHC and its 95% confidence interval (CI), calculated from beta and standard errors estimated by MaAsLin2 (square with solid line of 95% CI), and ANCOM-BC (circle with dotted line of 95% CI). Points and lines for fold changes and 95% CI were colored blue (elevated in PD) or red (reduced in PD).

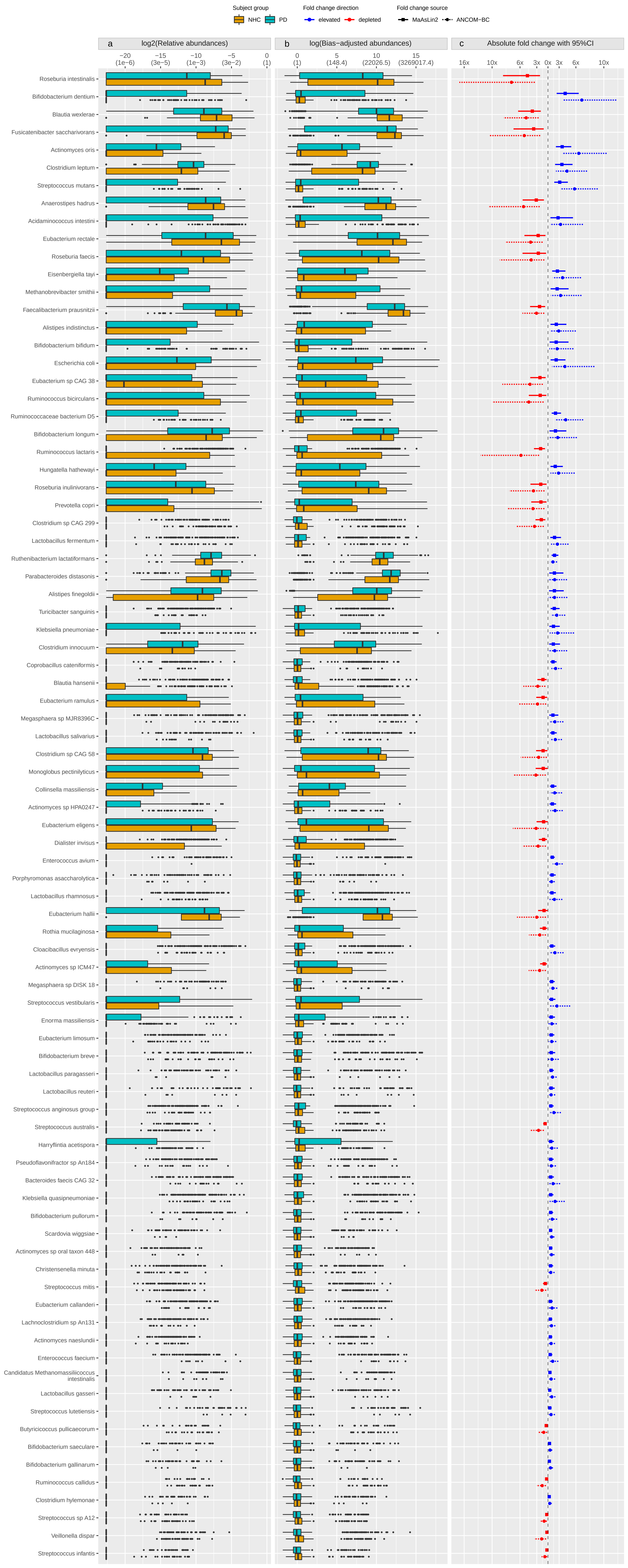

Subject group
NHC
PD
Fold change direction
elevated
depleted
Fold change source
MaAsLin2
ANCOM-BC
a
log2(Relative abundances)
−20 (1e−6)
−15 (3e−5)
−10 (1e−3)
−5 (3e−2)
0 (1)
b
log(Bias−adjusted abundances)
0 (1)
5 (148.4)
10 (22026.5)
15 (3269017.4)
c
Absolute fold change with 95%CI
16x
10x
6x
3x
0x
3x
6x
10x
Roseburia intestinalis
Bifidobacterium dentium
Blautia wexlerae
Fusicatenibacter saccharivorans
Actinomyces oris
Clostridium leptum
Streptococcus mutans
Anaerostipes hadrus
Acidaminococcus intestini
Eubacterium rectale
Roseburia faecis
Eisenbergiella tayi
Methanobrevibacter smithii
Faecalibacterium prausnitzii
Alistipes indistinctus
Bifidobacterium bifidum
Escherichia coli
Eubacterium sp CAG 38
Ruminococcus bicirculans
Ruminococcaceae bacterium D5
Bifidobacterium longum
Ruminococcus lactaris
Hungatella hathewayi
Roseburia inulinivorans
Prevotella copri
Clostridium sp CAG 299
Lactobacillus fermentum
Ruthenibacterium lactatiformans
Parabacteroides distasonis
Alistipes finegoldii
Turicibacter sanguinis
Klebsiella pneumoniae
Clostridium innocuum
Coprobacillus cateniformis
Blautia hansenii
Eubacterium ramulus
Megasphaera sp MJR8396C
Lactobacillus salivarius
Clostridium sp CAG 58
Monoglobus pectinilyticus
Collinsella massiliensis
Actinomyces sp HPA0247
Eubacterium eligens
Dialister invisus
Enterococcus avium
Porphyromonas asaccharolytica
Lactobacillus rhamnosus
Eubacterium hallii
Rothia mucilaginosa
Cloacibacillus evryensis
Actinomyces sp ICM47
Megasphaera sp DISK 18
Streptococcus vestibularis
Enorma massiliensis
Eubacterium limosum
Bifidobacterium breve
Lactobacillus paragasseri
Lactobacillus reuteri
Streptococcus anginosus group
Streptococcus australis
Harryflintia acetispora
Pseudoflavonifractor sp An184
Bacteroides faecis CAG 32
Klebsiella quasipneumoniae
Bifidobacterium pullorum
Scardovia wiggsiae
Actinomyces sp oral taxon 448
Christensenella minuta
Streptococcus mitis
Eubacterium callanderi
Lachnoclostridium sp An131
Actinomyces naeslundii
Enterococcus faecium
Candidatus Methanomassiliicoccus intestinalis
Lactobacillus gasseri
Streptococcus lutetiensis
Butyricicoccus pullicaecorum
Bifidobacterium saeculare
Bifidobacterium gallinarum
Ruminococcus callidus
Clostridium hylemonae
Streptococcus sp A12
Veillonella dispar
Streptococcus infantis

**Supplementary Fig. 4. Species correlation networks for PD and NHC.** All species that were detected were tested for pair-wise correlation in PD or NHC metagenomes separately. Correlation networks were constructed including species with a correlation of $|r| > 0.2$ and permuted P-value $< 0.05$ with at least one other species. Clusters of species within networks were defined by Louvain algorithm and were randomly assigned a color and a number. Each circle (node) denotes a species and the curved lines (edges) connect correlated species. 26 clusters were defined in the PD network and 20 in the NHC network. PD-associated species mapped to 12 clusters in the PD network (mainly in clusters 2, 8, 10 and 13), and 11 clusters in the NHC network (mainly in 2, 3, and 8). 244 of 719 species did not map to a cluster in either PD or NHC networks as they had no correlation with another species at $|r| > 0.2$ and $P < 0.05$, including 2 PD-associated species, one of which was elevated in PD and the other reduced. **(a)** PD metagenome network with algorithmically defined clusters of correlated species. **(b)** PD metagenome network with species that emerged from MWAS as having significantly higher abundance in PD than NHC are colored blue, and those that had significantly lower abundance in PD than NHC colored red. **(c)** NHC metagenome network with algorithmically defined clusters of correlated species. As colors and numbers were assigned randomly, they do not correspond to colors and numbers in PD network. **(d)** NHC metagenome network with species that emerged from MWAS as having significantly higher abundance in PD than NHC colored blue, and those that had significantly lower abundance in PD than NHC colored red.

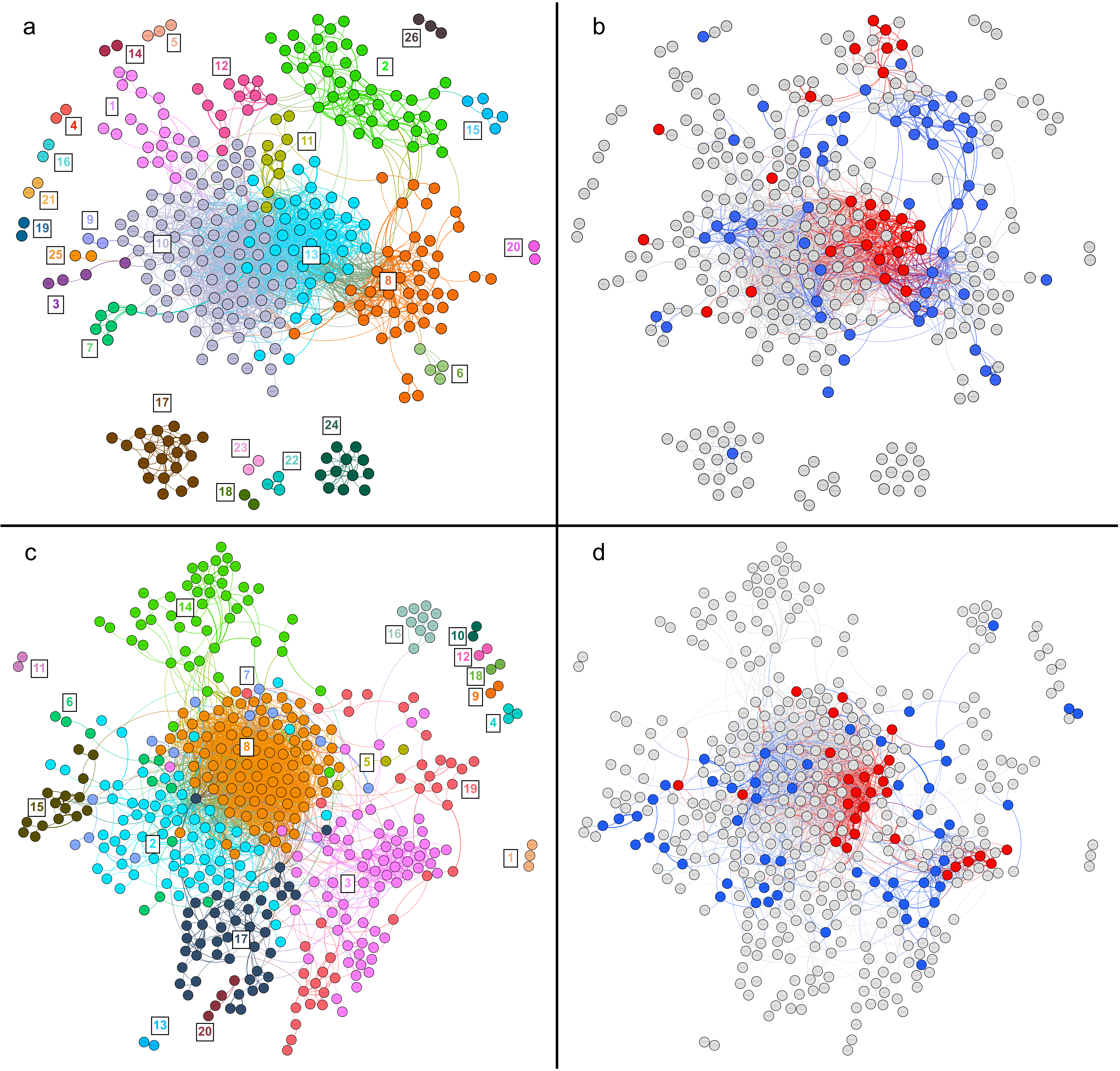
a
b
c
d
